# Linking somatic variants to altered methylation profiles in nanopore-derived breast and pancreatic cancer genomes using unduliner

**DOI:** 10.64898/2026.10.08.757715

**Authors:** Halimat C. Atanda, Adam D. Ewing

**Affiliations:** Mater Research Institute - University of Queensland, Brisbane, Australia

## Abstract

Research shows that genetic variation directly influences DNA methylation, and this association has generally been explored on germline SNPs and at a population level. The presence of genomic variation and base modification on sequence reads generated from a technology like nanopore sequencing offers an opportunity to link disease-specific variants to methylation changes in individual genomes. Here, we present unduliner, a framework with an embedded pretrained convolutional neural network model, that predicts somatic variants, single-nucleotide (SNVs) and structural (SVs), associated with methylation pattern changes in cancer genomes. We trained the model (accuracy = 0.803 on validation; 0.897 on held-out test) using phased sequence reads, single-nucleotide germline variants, per-base methylation probability, and variant proximity or presence in haplotype-specific differentially methylated regions (DMRs). Unduliner was applied to subline-specific variants from MCF7 breast cancer cell lines as proof-of-concept, and we demonstrated its ability to identify variant-stratified methylation differences independent of haplotype-based methylation differences. In a pancreatic ductal adenocarcinoma (PDAC) sample, we identified coding and non-coding SNVs and SVs associated with tumour-normal DMRs, allele-specific methylation, and intratumour DMRs. These observations demonstrate that integrating long-read methylation and variant information can provide additional interpretative context for both coding and non-coding genomic variation beyond the conventional variant impact prediction tools. The results from unduliner include variant information, size and direction of methylation change with respect to the variant (increase or decrease), and genomic location of methylation change. Unduliner is publicly available on GitHub at https://github.com/Chisomgold/unduliner.

## Introduction

DNA CpG methylation is the most observed and most studied epigenetic modification in the human genome (Moore et al., 2013; Sergeeva et al., 2023). It is a central component of genetic regulation, playing roles in gene expression, chromatin organisation and cellular identity. Whole methylome studies have identified global demethylation and focal hypermethylation at certain genomic regions in tumours compared to healthy genomes, making aberrant methylation an undisputed hallmark of cancers. Therefore, several approaches have been adopted to study, understand, and apply methylation markers to cancer management. Alongside methylation changes, cancer genomes also accumulate genetic variants, including single-nucleotide variants (SNVs) and structural variants (SVs), spread across the coding and non-coding regions (Campbell et al., 2020; Vogelstein et al., 2013) with potential regulatory consequences. While genetic variants and aberrant methylation have been studied and characterised individually, understanding the range of their interactions remains underexplored.

A growing body of evidence (Liang & Weisenberger, 2017; Nebbioso et al., 2018; Oliva, 2023; Stefansson et al., 2024) suggests that genetic variation can directly influence DNA methylation levels intra- and inter-chromosomally. Association studies have linked SNPs to individual CpG loci changes across populations in what is called methylation quantitative trait loci (meQTLs). In addition, studies have explored patterns, defined as epialleles or methylation haplotype blocks or allele-specific methylation (ASM), including the distribution of methylation across regions or co-occurrence of hypo- or hyper-methylation in adjacent bases. However, many of these studies have been driven by technologies, like microarrays and bisulfite sequencing, with limited ability to link specific variants to precise genomic regions over which methylation changes occur (Gaunt et al., 2016; Seoighe et al., 2018; Wang et al., 2021; Yuen et al., 2021). As a result, these studies fall short in different contexts, such as being limited to predefined CpG sites and the inability to resolve intratumour variant-methylation association for individual genomes.

Nanopore technology generates rich, high-dimensional data that can overcome these limitations (van Dijk et al., 2018; Yuen et al., 2021). The simultaneous detection of genetic variants and base-resolved DNA methylation states from single molecules and outputs spanning large genomic regions and supporting phasing creates the opportunity to directly link sequence variants to changes in methylation patterns within the same reads. The emergence of long read sequencing gave rise to new computational methods for bioinformatics tasks such as basecalling, variant analysis, and genome assembly, but none that fully leverage its capabilities to identify variant-associated methylated changes.

Here, we present a computational framework, unduliner, that identifies variant-associated DNA methylation changes from nanopore sequencing data. The method is designed to capture associations arising from both single-nucleotide variants and structural variants using methylation-aware (5mC) sequence reads and provide interpretable outputs on differentially methylated regions (DMRs) that will facilitate variant prioritisation and downstream biological analysis. We apply the method to subline-specific variants obtained from two sublines of the MCF7 breast cancer cell line (Atanda & Ewing, 2026a; Cheetham et al., 2022) as a proof-of-principle system and report the association of these variants with local methylation changes across distinct regions on the genome. We also demonstrate its utility using genomic data from a pancreatic ductal adenocarcinoma (PDAC) sample (McDaniel et al., 2025; Wagner et al., 2026).

## Results

### Unduliner prioritises variant-associated methylation signals

The final model architecture in unduliner comprised 4 convolutional layers (Fig. 1). On the validation set, the model achieved an accuracy of 0.803 (F1 = 0.806, MCC = 0.617).

**Figure 1.**
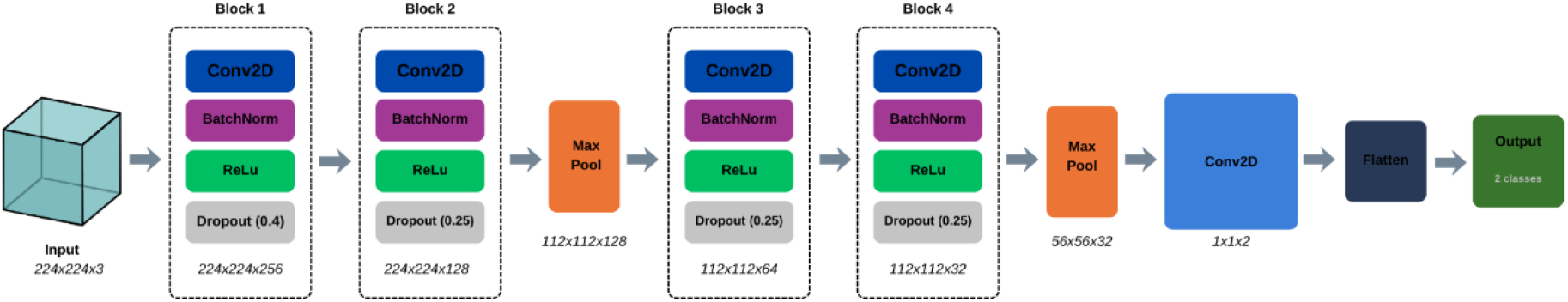
Abstract representation of unduliner’s CNN architecture

Performance on the held-out test partition improved further with an accuracy of 0.897 (F1 = 0.895, MCC = 0.791), indicating stable generalisation within the distribution.

Given the limited number of known associations between somatic variants and methylation changes, our training data comprised germline variants associated with haplotype-specific differential methylation (see Methods section). To assess performance beyond the training data, we applied the model to phased variants on chromosome 4 (n=6543) of the MCF7 ATCC subline. After excluding variants lacking read support (n = 178), 6365 variants were evaluated. As an approximate baseline, we used DSS-derived DMRs from haplotype-based comparisons. Variants were grouped based on their proximity (within ±10 kbp) to a DMR (areaStat > 200), mirroring the criteria used during model development. However, this framework is an indirect approximation constrained by a fixed genomic window. In contrast, our framework directly compares genotype-stratified read groups and is not constrained by predefined window sizes. Given that ONT reads often exceed 20 kbp in length, methylation effects extending beyond this window will not be captured by the DMR-based definition, restricting us from providing the conventional comparison performance metrics on this dataset.

Nevertheless, overlap with DSS-supported variants was enriched among the highest-ranked predictions, and visual inspection of the top-ranked DMRs (Supplementary Table 1) using methylartist --splitvar option (Supplementary Figure 1) confirmed clear subline-specific variant-stratified methylation differences independent of haplotype-based methylation differences. These examples show that high-confidence predictions correspond to visually evident methylation shifts linked to variants.

Importantly, unduliner outputs are ranked by statistical significance of the estimated methylation differences (FDR<0.05 default), and enable users to apply stricter thresholds and further prioritise high-confidence candidates. These results demonstrate that unduliner effectively identifies and prioritises variant-associated methylation differences in an interpretable manner that aligns with the underlying structure of long read sequence data.

### Unduliner workflow and design

Unduliner is a read-level framework for predicting somatic variant–associated methylation differences from ONT data (Fig 2). It has two modes of operation: the default expects a variant call format (VCF) file containing SNVs as one of the inputs, and SV analysis requires a specified option (--sv). Conceptually, unduliner works on image-encoded data created as described in the model training and development section (methods), where the columns represent aligned genomic sites, and each row represents a read. It accepts long-read alignment files with modification tags (MM/ML) as input alongside a standard VCF file containing the variants of interest. First, it creates bam file pairs for each variant, all of which are indexed and then arrays of 1,0,-1 depending on the extracted methylation probability, which is then used for image generation. The provided pretrained model makes a prediction for each image.

**Figure 2.**
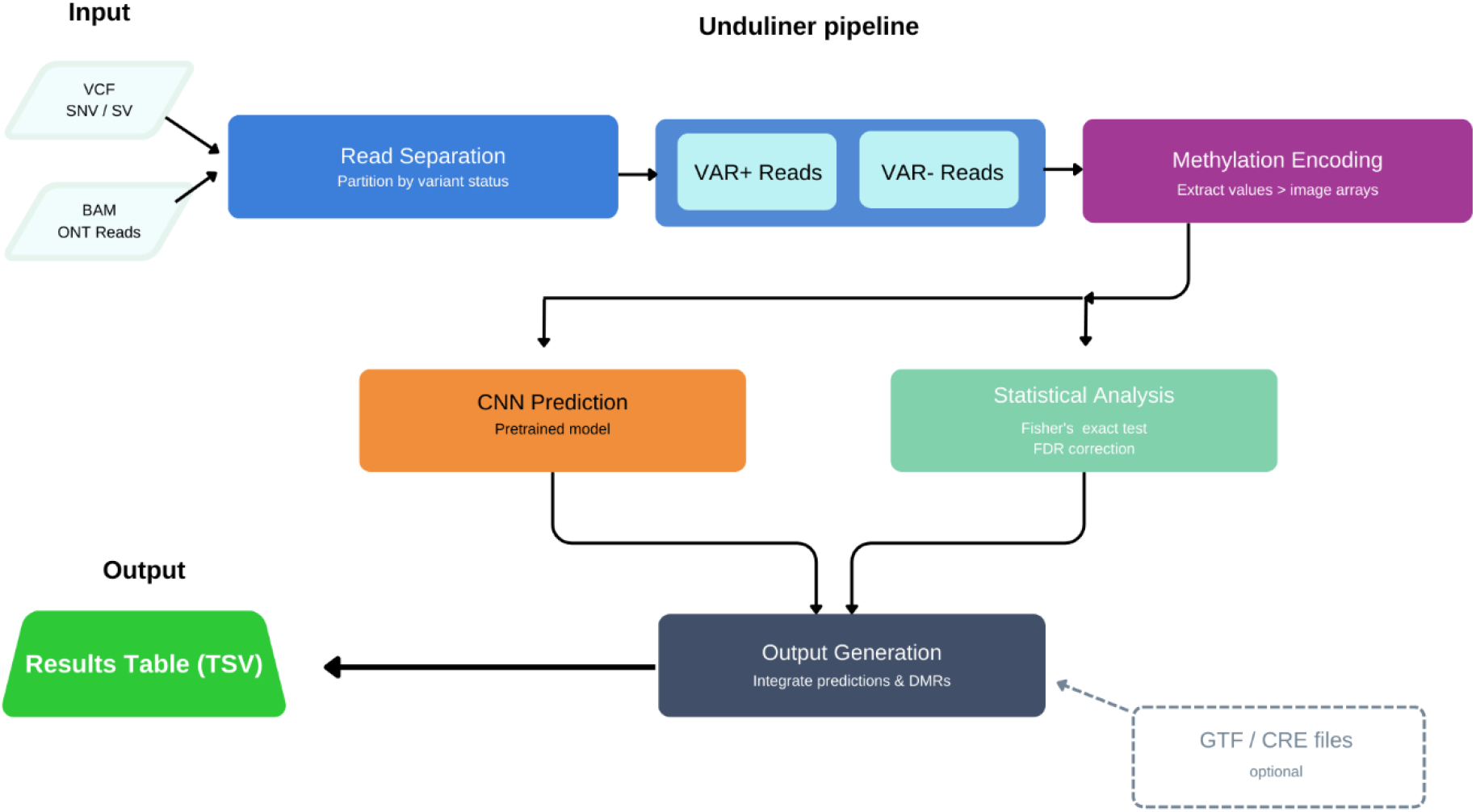
A high-level summary of the steps involved in unduliner analysis

To identify specific DMRs after prediction, unduliner compares methylation calls between reads carrying the variant allele and reads carrying the reference allele at aligned CpG positions. At each CpG site *i*, methylation status is encoded as methylated (+1) or unmethylated (−1). Sites with insufficient coverage in either group are excluded; by default, a minimum coverage of *C* reads per group is required (*C* = 3).

For each retained CpG site, the methylation difference (Δ_i_) is computed as the difference in methylation fractions between the two groups:

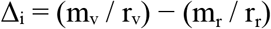

where m_v_ and m_r_ denote the number of methylated reads for the variant and reference alleles, respectively, and r_v_ and r_r_ denote the total number of reads with the variant and the reference base, respectively.

CpG sites are classified as differentially methylated or similar based on the distribution of Δ_i_ values. When multiple CpG sites are present on the reads being assessed, a two-component Gaussian mixture model is fitted to Δ_i_ values, with one component representing sites with minimal methylation differences and the other representing differential sites. In rare cases where fewer than two CpG sites are present on the reads, classification is instead based on a fixed absolute difference threshold |Δ_i_| > τ (τ = 0.2 by default). Adjacent differentially methylated CpGs are grouped into candidate DMRs if they are separated by no more than *D* base pairs (*D* = 50) and if the region contained at least *N* CpGs (*N* = 3). Regions shorter than *L* base pairs (*L* = 50) are discarded. Nearby DMRs separated by gaps of at most *G* base pairs (*G* = 500) are merged unless there is an overlap with non-differentially methylated CpGs. For each final DMR, methylated and unmethylated read counts are summed across all CpGs in the region, and statistical significance is assessed using Fisher’s exact test. Resulting *p*-values are corrected for multiple testing using the Benjamini–Hochberg false discovery rate procedure. For each tested variant, a maximum of 5 identified DMRs along with their corresponding *p*-values and quantified methylation differences are provided. All defined variables are user-adjustable.

In cases where variants are not supported by any reads in the sample bam file, that is, artefacts and/or technical errors, or variants are completely homozygous (r_r_ =0), unduliner does not process such variants and chromosomal information for each variant is stored and saved into a separate text file (example in Supplementary document). The major implication of this feature is that false positive variants have zero chances of being attributed to methylation changes. Unduliner also provides options to annotate associated methylated regions with relevant genomic features (GTF file) and regulatory elements like promoters (BED-format file with no header). Analysis can be limited to a specific chromosome or to a specific region on a named chromosome, and its runtime is expectedly affected by the number of variants being analysed and by sequencing depth.

Unduliner outputs a tab-separated BED-like table containing variant information, number of reads supporting the alternate and reference alleles, the model’s prediction, associated regions with differential methylation ranked by corrected p-values, and effect size of the difference, where a negative value indicates the regions surrounding the reference base are hypermethylated compared to those around the variant and a positive value indicates otherwise.

### MCF7 subline-specific variants exhibit monoallelic and biallelic presence with different mechanisms of methylation alteration

We applied unduliner to previously generated nanopore sequencing data from MCF7 cells, comprising one high-depth sample per MCF7 subline (ATCC and ECACC), obtained by combining sequencing runs as previously described (Cheetham et al., 2022). And we used previously generated subline-specific SNVs and SVs (Atanda & Ewing, 2026a) as the variants of interest.

In the ATCC subline, unduliner identified 405 subline-specific SNV-associated DMRs overlapping 313 genes. Among these genes, 20 are present in the Cancer Gene Census record, including 3 associated with breast cancer. The observed associations primarily represent 2 mechanisms: variant-induced methylation alterations and variant-amplified methylation differences (Supplementary Table 2). The absence of haplotype-associated differences in the former group suggests that the observed differential methylation emerged specifically in association with the subline-specific variant, whereas variants in the latter group are more likely modulators that exacerbate existing haplotype-specific methylation states. The ECACC subline yielded 40 subline-specific SNVs associated with differential methylation at regions that overlapped 28 genes (Supplementary Table 3), none of which have been implicated in cancer.

Heterozygous somatic (or subline-specific) variants are typically expected to arise on a single haplotype and result in allele-specific effects (Heyne et al., 2023). However, our results show a different landscape, where variants associated with methylation changes are either allele-specific or biallelic. While some of these observations could be a consequence of incomplete or ambiguous phasing, they could also be biologically relevant as instances of biallelic germline and somatic mutations have been reported in tumour genomes (Demeulemeester et al., 2022).

In this case, where variants were confined to a single haplotype, methylation differences often pre-existed at the haplotype level and were amplified upon variant-based partitioning. For example, in the ATCC subline, an intronic SNV in GATA3 (chr10:8066630 G>A) shows substantial allele-specific differential methylation at the regions identified by unduliner (Fig 3a(i)), and this difference becomes even more pronounced when reads are stratified by variant status (Fig 3a(ii), Supplementary Figure 2).

**Figure 3.**
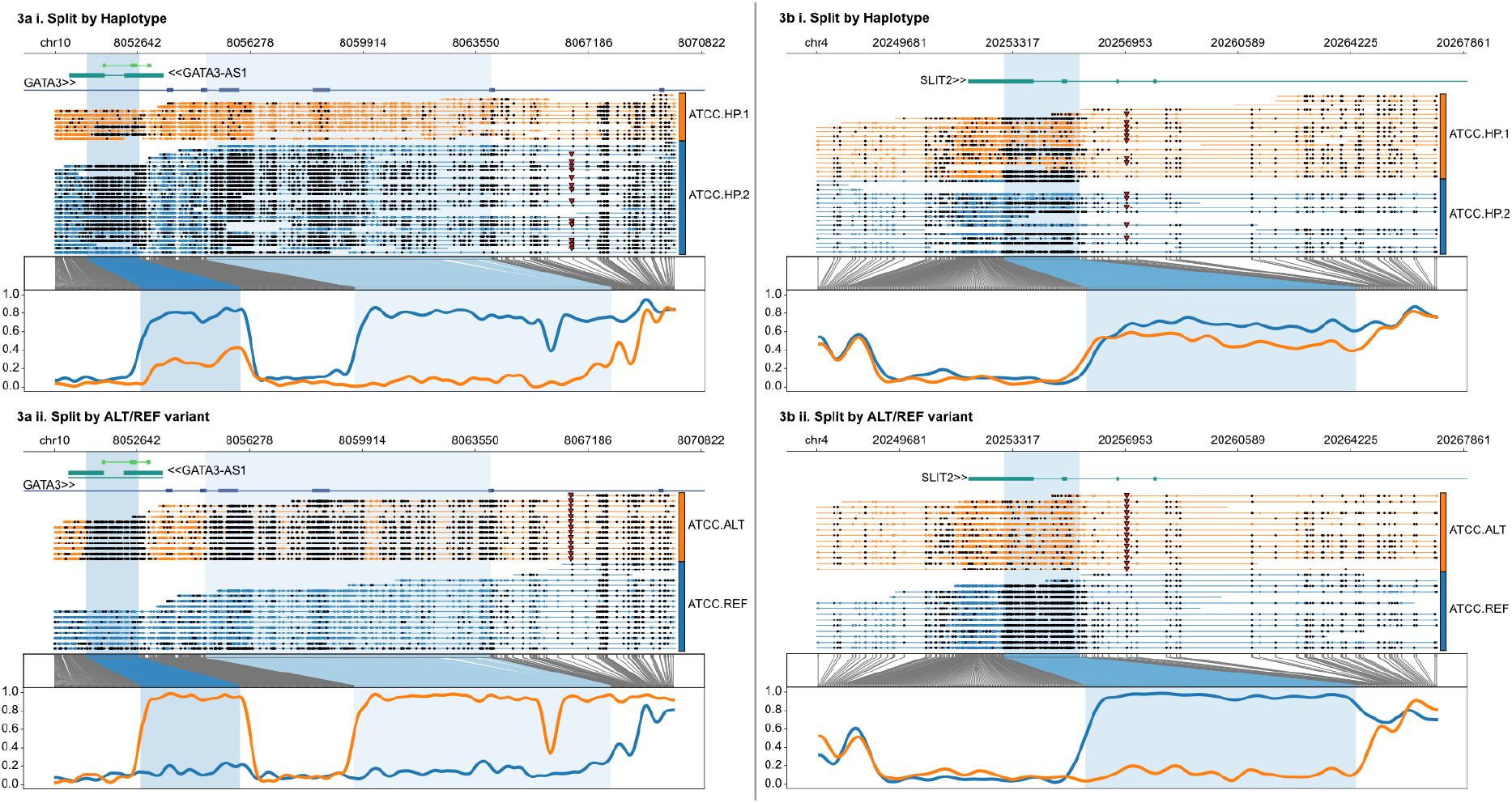
Example of “variant-amplified” (3ai&ii) and “variant-induced” (3bi&ii) differential methylation relative to haplotype-level methylation. The variants are marked by triangles.

In contrast, where variants are biallelic, they are often characterised by minimal allele-specific differential methylation at unduliner-identified regions which then show strong alteration upon variant stratification. An example in the ATCC subline is an intronic SNV in SLIT2 (chr4:20256997 G>A), where reference-supporting reads are nearly fully methylated while variant-supporting reads are largely unmethylated at the highlighted region (Fig 3b(i)(ii), Supplementary Figure 3). This DMR also overlaps active regulatory features, suggesting a shift towards increased regulatory accessibility.

Similar patterns (Supplementary Figures 4 & 5) were observed in the ECACC subline across distinct genomic contexts. These observations highlight the ability of a read-based approach to separate haplotype-driven epigenetic effects from variant-driven ones.

For structural variants, we identified 518 subline-specific SVs associated with DMRs in the ATCC subline and 16 in the ECACC subline (Supplementary Tables 4a&b). The ATCC DMRs overlapped 386 genes, including 13 cancer-associated genes (3 linked to breast cancer), whereas the ECACC DMRs overlapped 10 genes, with 2 being cancer-associated. Equivalent classification of SV—methylation effects into induced versus amplified relative to haplotype-level methylation was not feasible due to limitations in visually resolving several SV-associated read-split signals at the haplotype level, and in some cases, the genomic region of the SV was unphased. Instead, a few illustrative examples of variant-associated differential methylation are presented per subline (Fig 4).

**Figure 4.**
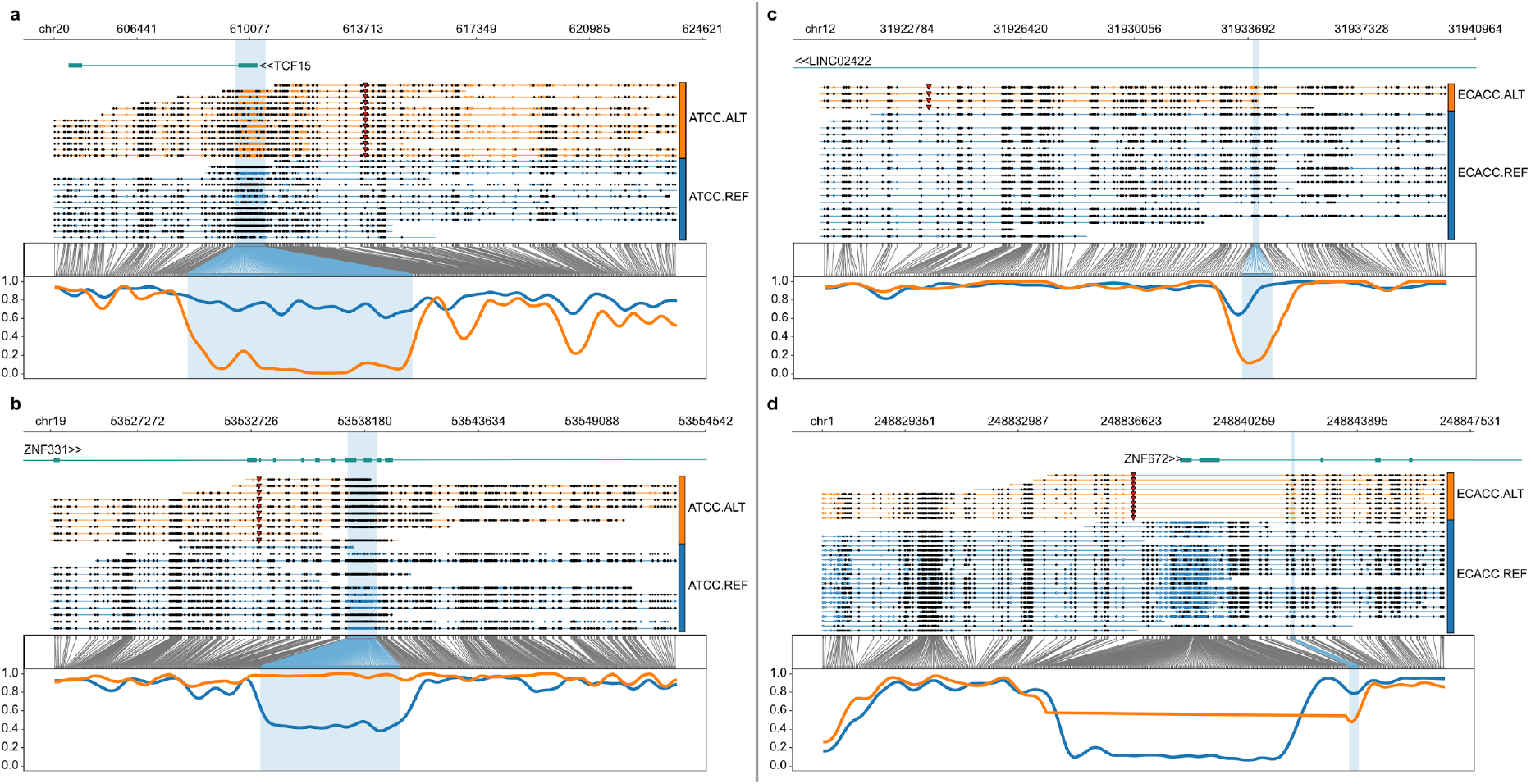
Methylartist plot of SVs and associated DMRs from the ATCC (left) and ECACC (right) sublines. a) Deletion on chr20 and associated DMR with hypomethylated variant reads overlapping regulatory regions on TCF15 gene. b) Insertion on chr19 and associated DMR with hypermethylated variant reads overlapping regulatory regions around ZNF331. c) Insertion on chr12 and associated DMR on LINC02422 with hypomethylated variant reads. d) Deletion on chr1 and associated DMR on ZNF672 with hypomethylated variant reads.

### Variant-associated methylation changes preferentially affect distal regulatory elements in MCF7

Applying unduliner to subline-specific variants revealed widespread associations between genetic variation and local methylation differences. These effects spanned a broad genomic range, with variant–DMR distances ranging from 2 bp to ∼90 kbp (Fig 5), with several regions overlapping the variants, indicating that genetic variation influences methylation within and beyond its immediate genomic context.

**Figure 5.**
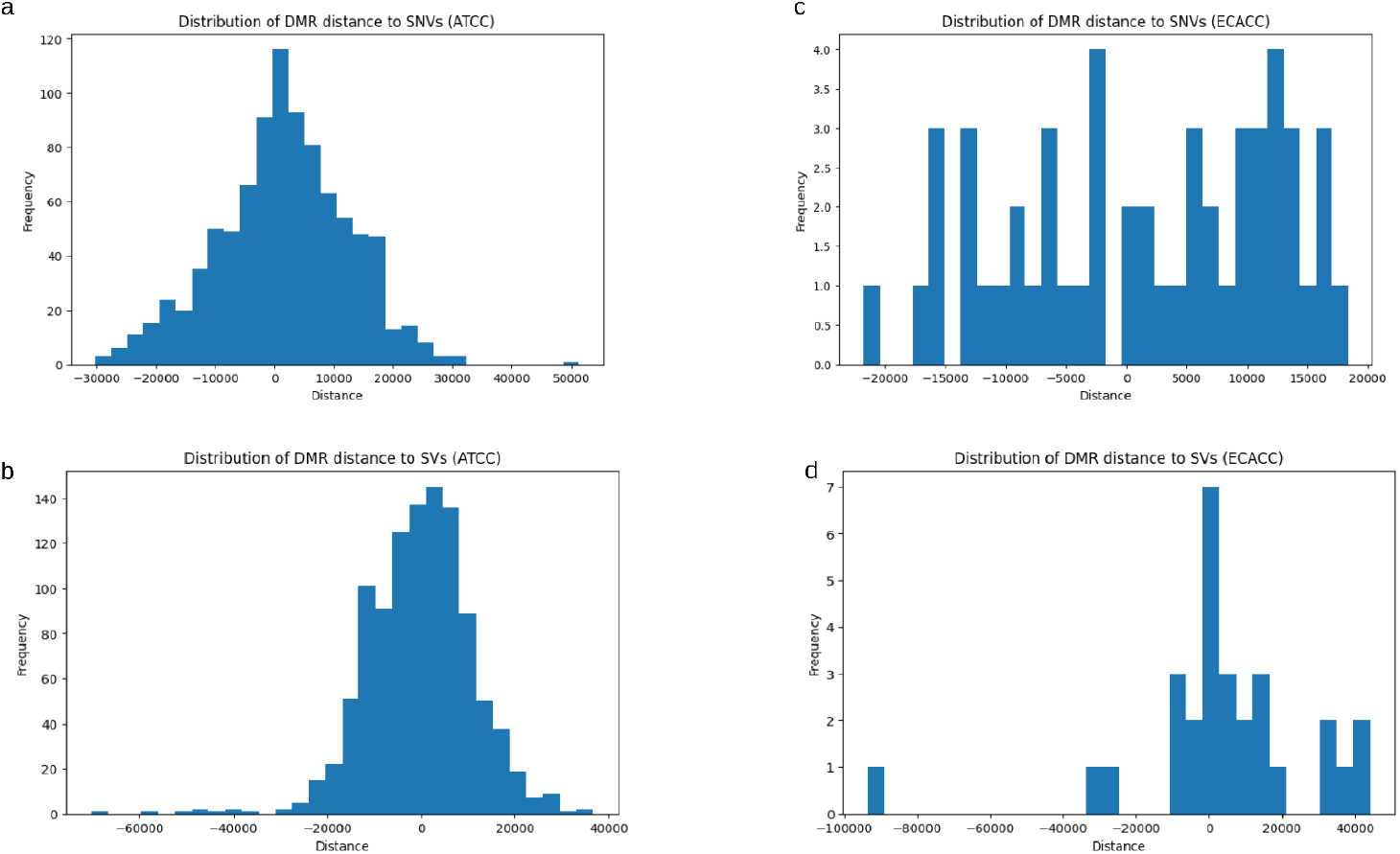
The distribution of the genomic range between variants and DMRs in the ATCC (a & b) and ECACC (c &d) sublines, where distance is the difference between the DMR end position and the variant position.

Consistent with this observation, variant-associated DMRs were significantly enriched in enhancer-like elements, based on annotations from ENCODE3 cCREs (Table 1). This enrichment supports a model in which genetic variation disturbs regulatory landscapes as well as coding sequences. It highlights the importance of noncoding variation in shaping epigenetic states. Also, given the distal nature of these variant—DMR relationships, the limited overlap of the variant-associated DMRs with known cancer-related genes underscores the limitations of gene-centric interpretation of genetic variation.

**Table 1.**
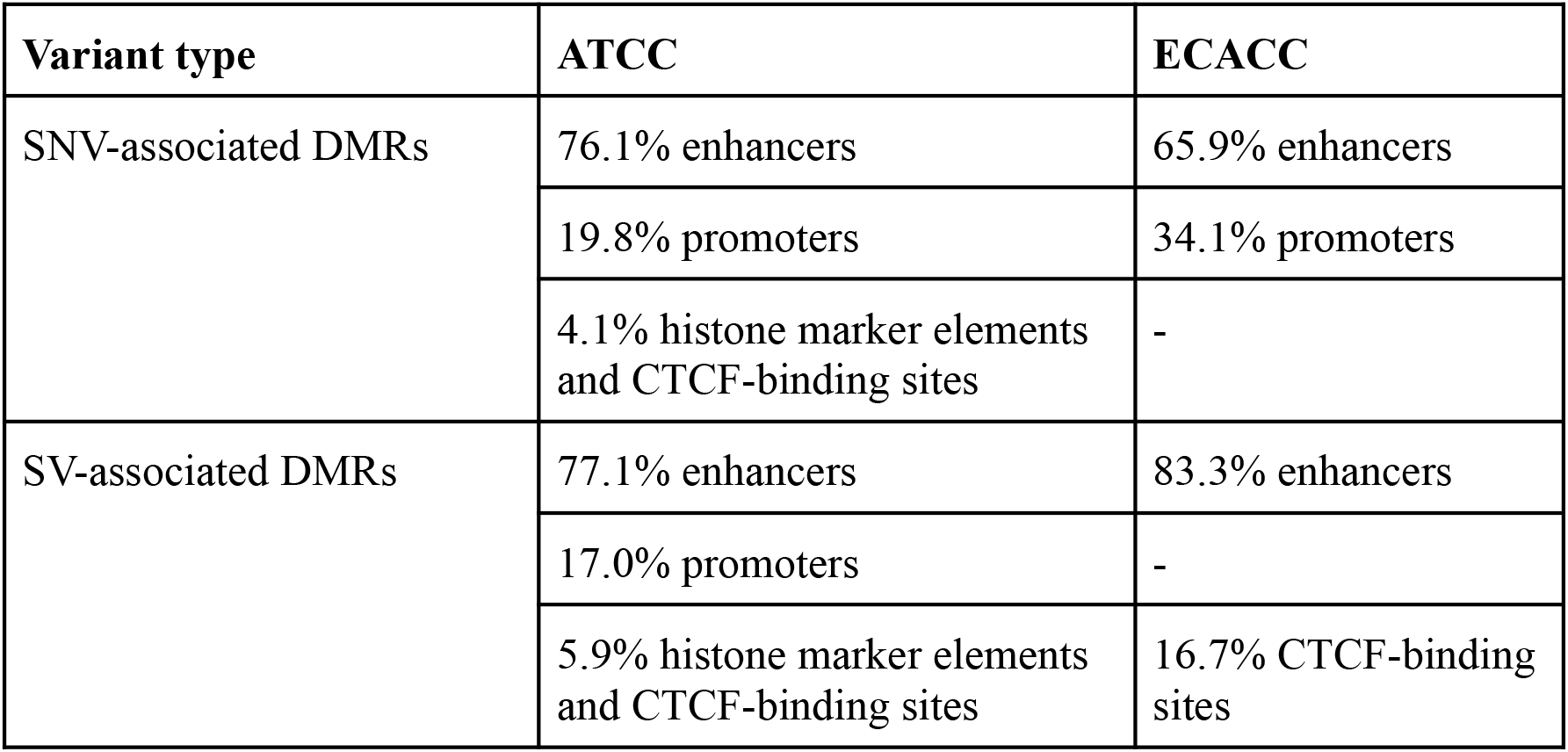
Enrichment of variant-associated DMRs with known regulatory elements.

### Variant-associated reads in MCF7 show a bias towards hypomethylation

We observed a clear directional bias, with variant-associated reads more frequently exhibiting hypomethylation compared to reference reads at identified loci in the context of both SNVs and SVs in both sublines (Table 2&3). This suggests that most of the observed altered methylation may preferentially disrupt methylation maintenance and contribute to regulatory activation.

**Table 2.** Summary table of SNV-associated DMRs for ATCC and ECACC sublines.

| **Metric** | **ATCC subline** | **ECACC subline** |
| --- | --- | --- |
| Total positive variants (after filtering) | 405 | 40 |
| Total variant-loci associations | 599 | 41 |
| Hypermethylated loci (%) | 30.0 | 24.4 |
| Hypomethylated loci (%) | 70.0 | 75.6 |
| Total overlapped genes | 313 | 28 |
| Total overlapped cancer-related genes | 20 | - |

**Table 3.** Summary of SV-associated DMRs for ATCC and ECACC sublines.

| **Metric** | **ATCC subline** | **ECACC subline** |
| --- | --- | --- |
| Total positive variants | 518 | 16 |
| Total variant-loci associations | 1054 | 29 |
| Hypermethylated loci (%) | 36.4 | 31.0 |
| Hypomethylated loci (%) | 63.6 | 69.0 |
| Total overlapped genes | 386 | 10 |
| Total overlapped cancer-related genes | 13 | 2 |

### Unduliner identifies variant—methylation associations in primary PDAC genome

We identified 9995 tumour-specific SNVs and 290 SVs. At default setting of unduliner (FDR < 0.05 and |methylation-difference| >0.2), approximately 73% of assessable, that is, excluding false positive variants, tumour-specific SNVs (6910/9415) and 43% SVs (106/242) were labelled as positively associated with methylation changes (Supplementary Table 5a&b), with variant-to-DMR distances ranging from 1 bp to ∼298 kbp (Supplementary Figures 6-9). While at face value, this seems like an unexpectedly high number of candidates, it can be explained partly by the default settings and partly by our observation in some instances where multiple closely located somatic variants covered by the same reads were associated with the same region of differential methylation.

Similar to the observation in the MCF7 analyses, positive variants were either limited to one haplotype or shared across both, and associated DMRs often showed different methylation patterns in variant-stratified groups compared to haplotype-stratified groups (Supplementary Figures 10&11). We also observed instances of variant-linked methylation differences that could explain tumour-normal differential methylation (Supplementary Figures 12-16), intratumour differential methylation absent in tumour-normal comparison (Supplementary Figure 17), and variant—methylation associations linked with allele-specific differential methylation (Supplementary Figures 18 &19).

From a regulatory context, 44% (12776/29030) of unduliner region hits in SNVs overlapped regulatory elements, and 80% of these were enhancers, 12% were promoters, and the remainder comprised H3K4me3 regions and CTCF binding sites. For SV-associated regions, 11% (39/351) overlapped regulatory elements, with enhancers being the most prominent.

Repetitive element overlap was assessed using UCSC repeat annotations. Among SNV-associated regions, 81% overlapped repeat elements: 43% SINEs, 23% LINEs, 15% LTRs, 10% microsatellites, and others, including retrotransposons. Among SV-associated regions, 85% (297/351) overlapped repeats: 43% SINEs: 17% satellites, 14% LINEs, and 9% microsatellites. The high representation of repeats in these variant-associated regions is consistent with the known susceptibility of repetitive sequences to methylation dysregulation in cancer genomes.

### Software

The core algorithm, functions and command-line front-end of unduliner is written in Python 3. External libraries include Pysam (https://github.com/pysam-developers/pysam) for bam and VCF parsing and Pytorch (Ansel et al., 2024) for the neural network implementation. The neural network was trained on a GPU, but using unduliner does not require one.

### Code availability and usage

Unduliner is open source, documented, and available at GitHub under the MIT license (https://github.com/Chisomgold/unduliner) as well as on Zenodo (Atanda & Ewing, 2026b). Provided installation options include repository cloning and pip. Data for a test run are available at https://github.com/Chisomgold/unduliner/tree/main/testdata.

## Discussion

Interpreting the functional or regulatory consequences of genetic variation is a central challenge in genomics and biomedical research (Foissac et al., 2012), particularly for noncoding regions. Many existing variant impact prediction or interpretation frameworks prioritise coding sequences (Richards et al., 2015) or rely on population-level associations, such as expression quantitative trait loci (eQTLs) (Ardlie et al., 2015) and methylation quantitative trait loci (meQTL) (Smith et al., 2014). While these approaches have been valuable in linking genetic variations to molecular phenotypes, they are inherently limited in their ability to resolve variant effects at single-sample resolution.

We present a read-based framework, unduliner, that directly links somatic or sample-specific variants to local DNA methylation changes using long-read nanopore sequencing data.

Leveraging the per-read co-occurrence of nucleotide variant and methylation information enables unduliner to identify variant-associated DMRs within individual samples, which aligns with the existing evidence that regulatory variation is highly context-specific and can be missed by aggregate analysis (Ardlie et al., 2015; Foissac et al., 2012; Liu et al., 2025). This sample-specific feature is fitting for scenarios where intratumour heterogeneity and regulatory variations require resolution without population-scale data. In addition, it effectively assigns regulatory association to structural variants, despite their reputation as being challenging to study due to their size and complexity. Finally, and perhaps most strikingly, the field of noncoding variant effect prediction is rapidly growing, and DNA methylation as a feature or an output is absent from some of the prominent tools in this space. For instance, PromoterAI (Jaganathan et al., 2025) predicts the effects of noncoding promoter variants on gene expression after being pre-trained to predict histone modifications, transcription factor binding, chromatin accessibility, and strand-specific gene expression but not methylation. Similarly, AlphaGenome (Avsec et al., 2026), the most comprehensive sequence-to-function model available as at 2026, predicts variant effects across several modalities, including gene expression, histone modifications, splice site usage and splice junction coordinates, but excludes methylation, which was highlighted as a direction for future improvements. For cancer genomics, where methylation reprogramming is one of the main features of malignant states, the omission of methylation from these tools is difficult to justify. AlphaGenome’s developers also highlighted that their training data and evaluations were heavily focused on protein-coding genes, and unduliner does not carry this constraint, as its training data was selected without any bias towards genomic annotation or filtering by coding status or regulatory overlap. While unduliner does not compete with these tools in scope or computational sophistication, it addresses a dimension of variant interactions, across coding and noncoding regions, that they do not explore.

Application of unduliner to breast cancer sublines and a PDAC sample demonstrated widespread associations between subline-specific variants and local methylation changes in regions strongly enriched in regulatory and repeat elements. We also show that variant-based methylation differences are not repetitive of, and cannot be reduced to, haplotype-linked methylation. In the context of variant types, distance to target (cis-associations), and genomic annotations of affected regions, our findings are relatively in line with previous similar studies focused on meQTL characterisation on a population level using short read technology (Banovich et al., 2014; Smith et al., 2014; Villicaña et al., 2023).

Despite unduliner’s ability to identify high-confidence associations between variants and methylation changes, it does not establish causality or downstream consequences. This will require integration with complementary omics data types, including RNA sequencing or chromatin accessibility assays. Also, the default thresholds in unduliner is permissive to reduce the risk of missing low-coverage regions and could result in more reported hits, as shown in the PDAC analyses; however, this can be managed by selecting the highest-ranked association, as they consistently exhibit visually validated differential methylation patterns. A further limitation is the absence of a comparable tool against which to benchmark unduliner’s performance. Benchmarking would typically involve running two or more tools on the same input data and comparing outputs against a reference or each other. The closest existing tool to unduliner, is DAMEfinder (Orjuela et al., 2020), which identifies allele-specific methylation differences from bisulfite sequencing data, a fundamentally different data type to direct read methylation state data. A meaningful comparison would require matched bisulfite and nanopore sequencing of the same sample while accounting for sequencing technology differences, and no such dataset was available for the samples analysed here. As corresponding datasets or similar tools across multiple platforms are available, a formal comparison will be possible.

In summary, unduliner provides a novel framework for single-sample read-based interpretations of genetic variation in a regulatory context. We expect that it will support a more comprehensive understanding of how genetic variation shapes methylation, and ultimately, cellular phenotype and disease susceptibility. The utility of unduliner has been demonstrated in cancer genomes; however, its underlying algorithm makes it applicable to identifying somatic variant—methylation associations in non-cancer genetic conditions.

## Method

### Data generation and encoding for model development

We obtained nanopore genome data of the ATCC MCF7 breast cancer cell line from Bioproject PRJNA748257 (Cheetham et al., 2022). Specifically, we used phased germline single-nucleotide variants (SNVs) from all available chromosomes except chromosome 4, which was set aside for independent testing, phased sequence alignment reads with whatshap (Martin et al., 2016), and haplotype-specific DMRs called with Dispersion Shrinkage for Sequencing data (DSS) Bioconductor tool (Wu et al., 2015). We defined variants within or in proximity (+/-10kbp) to identified DMRs (DSS areastat>200) as associated with local methylation pattern changes and classified these as positive. Variants not within 10kbp of a DMR were classified as negative. To minimise redundancy from clustering of germline variants that share proximity to a given DMR and exhibit correlated methylation profiles even when reads are separated by genotype, we implemented a distance-based filtering which enforced a minimum inter-variant distance of 20kbp when selecting variants for the model development.

We created bam file pairs per variant site, such that each site had a “with-variant” (reads supporting the alternate allele/variant) and “without-variant” (reads supporting reference allele) group, where the positive pair would have difference in methylation in at least one region (Fig 6a) and the negative pair would have no difference in methylation levels or non-specific methylation differences (Fig 6b). These bam files were individually sorted and indexed. We confirmed the correctness of this grouping strategy by randomly and manually inspecting some sites on Integrative Genome Viewer (IGV) and with methylartist (Cheetham et al., 2022).

**Figure 6.**
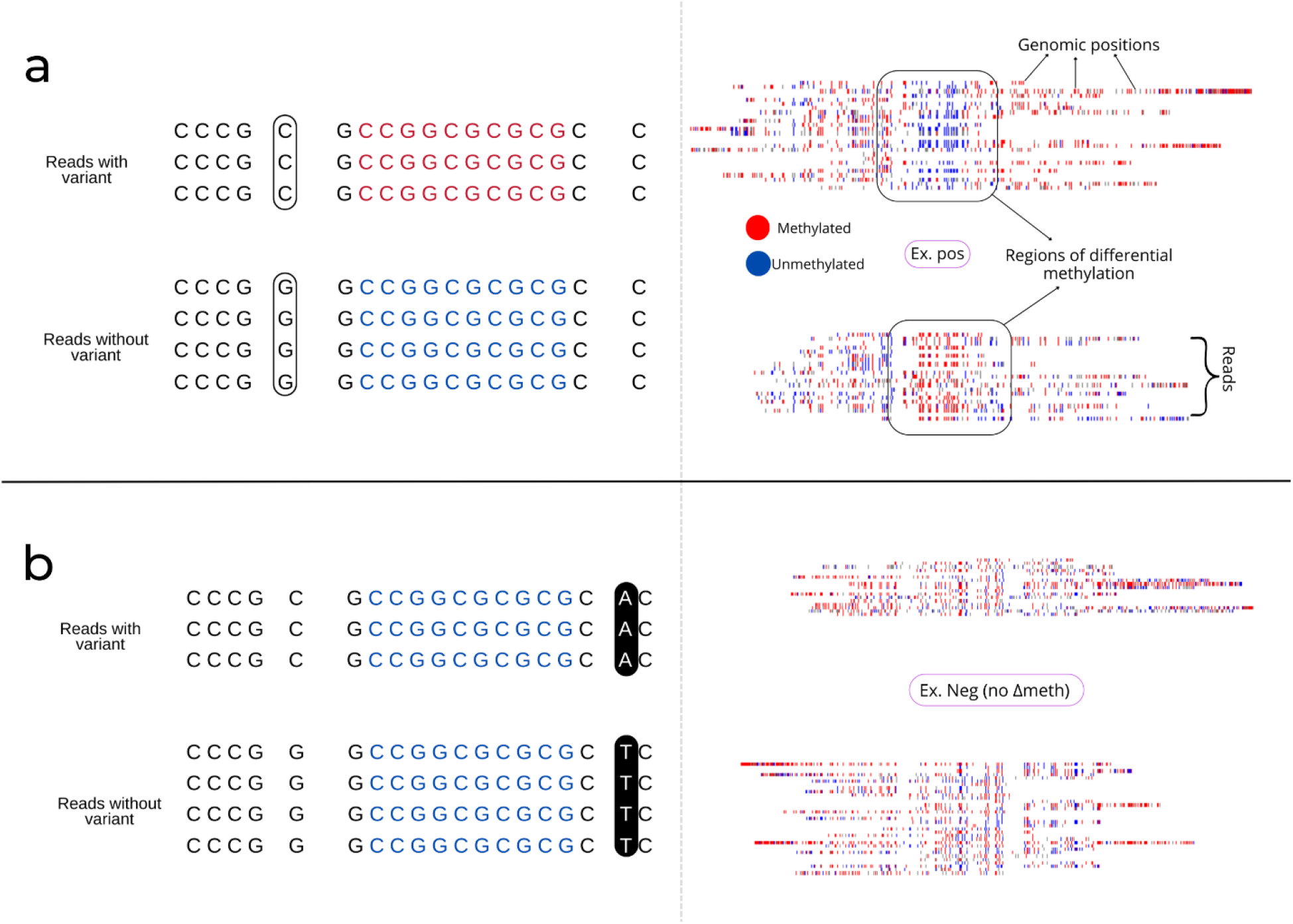
Side-by-side example of sequence read data encoding in a simplified illustration (left) and actual encoded image (right) showing read separation based on presence/absence of variant. (a) Separation of reads based on G/C (ref/alt) bases with different methylation profiles (positive pair example). (b) Separation of reads based on T/A (ref/alt) bases with similar methylation profiles (negative pair example).

Then, a 2D array of 5mC methylation values on each read was generated for each bam file and used to encode input data for the CNN as 2D RGB images (Fig 6). The arrays for the bam file pair (with-variant and without-variant) were plotted as subplots within the same figure if there were at least 3 reads per reference and alternate base, and if the total reads for one array is at least 30% of the other. This criterion ensured each subplot in the figure had enough visual information from which the model can learn. We represented nucleotide bases, specifically Cytosines (C), on a sequence read as colours reflecting their methylation probability values, obtained using the modified_bases function from the Pysam package. We approached this representation as discrete “trinary” values of 1, 0, and -1, where 1 equals probability >0.8 or methylated coloured red, -1 equals probability <0.2 or unmethylated coloured blue, and 0 equals 0.2<=probability<=0.8 or ambiguous/uncertain coloured grey (https://github.com/Chisomgold/unduliner/blob/main/dev-scripts/image_generator.py). The choice of colours was influenced by the default colour grouping of methylation on IGV.

### Model architecture, development and evaluation

We used the Pytorch deep learning framework to implement a convolutional neural network (CNN) architecture consisting of an initial series of convolutional blocks, each comprising a 2D convolutional layer followed by batch normalisation, ReLU activation, and dropout layers. The dropout rate was set to 0.4 for the first convolutional layer and 0.25 for subsequent layers. The kernel size for the first convolutional layer is 5 × 5, while subsequent layers use a kernel size of 3 × 3. Padding was adjusted accordingly to preserve spatial dimensions. The number of filters (output channels) starts at 256 for the first convolutional layer. It is halved after each layer to reduce computational cost and increase efficiency while retaining extracted features. Max pooling layers with a kernel size of 2 × 2 and a stride of 2 are added after every two convolutional blocks to reduce the spatial dimensions progressively. The final layer produces predictions for the two output classes.

We used the Adaptive Moment Estimation (Adam) optimiser, which has been reported to be memory and computationally efficient (Alzubaidi et al., 2021). Multiple models were trained using varying hyperparameter value combinations, including learning rates (0.0003, 0.0001, 0.005, 0.001, and 0.003), number of layers (3, 4, and 5), and batch sizes (32, 64, and 128) for a maximum of 200 epochs set to stop when training and validation accuracies reach 99% or when loss consistently fails to improve (https://github.com/Chisomgold/unduliner/blob/main/dev-scripts/modeltrain.py). While computer vision tasks often require very deep CNNs, sometimes up to 100 layers, previous research applying deep learning have shown that genomics tasks so far perform well on fewer convolutional layers (<=5) (Zou et al., 2019).

We randomly assigned variants from specific chromosomes to different dataset partitions; specifically, chromosomes 1, 2, 5, 7, and 10 for training; chromosomes 6, 11, 13-17, and 20 for validation, and chromosomes 3, 8, 9 and 19 for testing (Supplementary Tables 6a&b).

While this does not allow for a defined percentage split, it avoids potential data leakage that may result from similar “chromosome-specific” methylation patterns being in training, validation and testing datasets if using randomisation and a percentage-specific approach, which could cause the models to memorise rather than learn patterns. The final training dataset was imbalanced, with a higher negative (∼2x) to positive ratio.

Since the data was imbalanced, in addition to common metrics for classification tasks like accuracy and F1 score, we also report Matthews Correlation Coefficient (MCC) (Alharbi & Rashid, 2022; Alzubaidi et al., 2021; Chicco & Jurman, 2020). Independent model and tool testing were performed on phased variants from chromosome 4. We assessed performance by manual inspection of the predictions and highlighted regions from unduliner and using DSS-calculated haplotype DMRs.

### Whole genome subline-specific variant analysis for MCF7 sublines

We applied unduliner to subline-specific single-nucleotide and structural variants of ATCC and ECACC sublines of MCF7 (doi: 10.6084/m9.figshare.31360360) (Atanda & Ewing, 2026a; Cheetham et al., 2022) to determine which ones were likely associated with methylation changes. Positively identified variants and associated DMRs were compared to haplotype-level methylation differences, and we retained variant-specific DMRs that differed from haplotype-level DMRs for further investigation.

### Pancreatic ductal adenocarcinoma (PDAC) data retrieval and analyses

We obtained ONT whole-genome sequencing data of PDAC and matched non-tumour samples from the publicly available HG008 dataset developed by the Genome in a Bottle Consortium (McDaniel et al., 2025; Wagner et al., 2026) (https://ftp-trace.ncbi.nlm.nih.gov/ReferenceSamples/giab/data_somatic/HG008/Liss_lab/). Three tumour sequence datasets from different ONT libraries were downloaded and merged using samtools. We used ClairS (v0.4.4) (Zheng et al., 2023) to identify single-nucleotide variants, and structural variants were identified using Sniffles2 (v2.0) (Smolka et al., 2024), with both default parameters and the --mosaic option enabled to increase sensitivity to subclonal events. Variants with the PASS filter served as inputs to identify variant-methylation associations using unduliner.

Germline SNVs were called from GRCh38-aligned Illumina whole-genome sequencing data (https://ftp-trace.ncbi.nlm.nih.gov/ReferenceSamples/giab/data_somatic/HG008/Liss_lab/BCM_Illumina-WGS_20240313/) using GATK HaplotypeCaller (McKenna et al., 2010).

Variants were filtered to retain high-confidence heterozygous SNVs following GATK filtering thresholds https://gatk.broadinstitute.org/hc/en-us/articles/360035531112--How-to-Filter-variants-either-with-VQSR-or-by-hard-filtering). Phasing was performed with WhatsHap (v2.8) (Martin et al., 2023) to generate chromosome-scale phased VCF files, and phased variants were subsequently used to generate haplotagged ONT tumour and normal BAM files

### Variant-associated DMR annotation and visualisation

Variant-associated DMRs were annotated with genomic, cis-regulatory, and repeat elements obtained from the UCSC Genome Browser. Visualisation was enabled by the “--splitvar” and “--variants” options in methylartist locus.

## Acknowledgements

We acknowledge the Translational Research Institute (TRI) for research space, equipment, and facilities. This study was funded by the Australian Department of Health Medical Research Future Fund (MRFF) (MRF1175457 to A.D.E.) and by support from the Mater Foundation. This work was also supported by resources provided by The University of Queensland Research Computing Centre’s Bunya supercomputer (https://dx.doi.org/10.48610/wf6c-qy55), with funding from The University of Queensland, Brisbane, Australia.

