## Supplementary document for "Linking somatic variants to altered methylation profiles in nanopore-derived breast and pancreatic cancer genomes using unduliner"

Skipped the following positions because at least one BAM was empty:

No ALT-supporting reads for:
chr1:2648564-2648565;INS
chr1:3181551-3181552;INS
chr1:13151675-13224056;DEL
chr1:248433263-248632254;INV
chr2:89753992-89753993;BND
chr2:90325904-90325905;INS
chr4:89186205-89186206;INS
chr5:34197342-70485214;DEL
chr6:32526018-32526019;INS
chr6:32533022-32533023;INS
chr6:32543760-32543761;INS
chr6:166288742-166288743;INS
chr7:60997060-60997061;INS
chr7:61288279-61288280;INS
chr7:61327787-61603197;DEL
chr7:158149039-158149040;INS
chr9:43444400-43445083;DEL
chr10:97251704-97251705;INS
chr11:18965197-18965198;INS
chr12:37342784-37342785;INS
chr12:121442779-121442780;BND
chr13:16254866-16254867;BND
chr15:20238638-21037298;INV
chr16:1229157-1247503;INV
chr16:32806692-32806693;INS
chr19:11208145-11208146;INS
chr19:38981211-38983523;INV
chr20:1607146-1607147;INS
chr20:30456076-30456077;BND
chr21:5330013-7265075;DEL
chr21:5677595-5677596;BND
chr22:12275587-12275588;BND
chr22:15608569-15608570;INS
chr22:16381351-16381352;INS
chr22:18734438-18734439;INS
chrY:10860493-10860557;DEL
chrY:11295511-11295512;INS

No REF-supporting reads for:
chr2:89753992-89753993;BND

chr20:30456076-30456077;BND
chr21:5677595-5677596;BND
chr22:12275587-12275588;BND
