## Supplementary Figures for "Linking somatic variants to altered methylation profiles in nanopore-derived breast and pancreatic cancer genomes using unduliner"

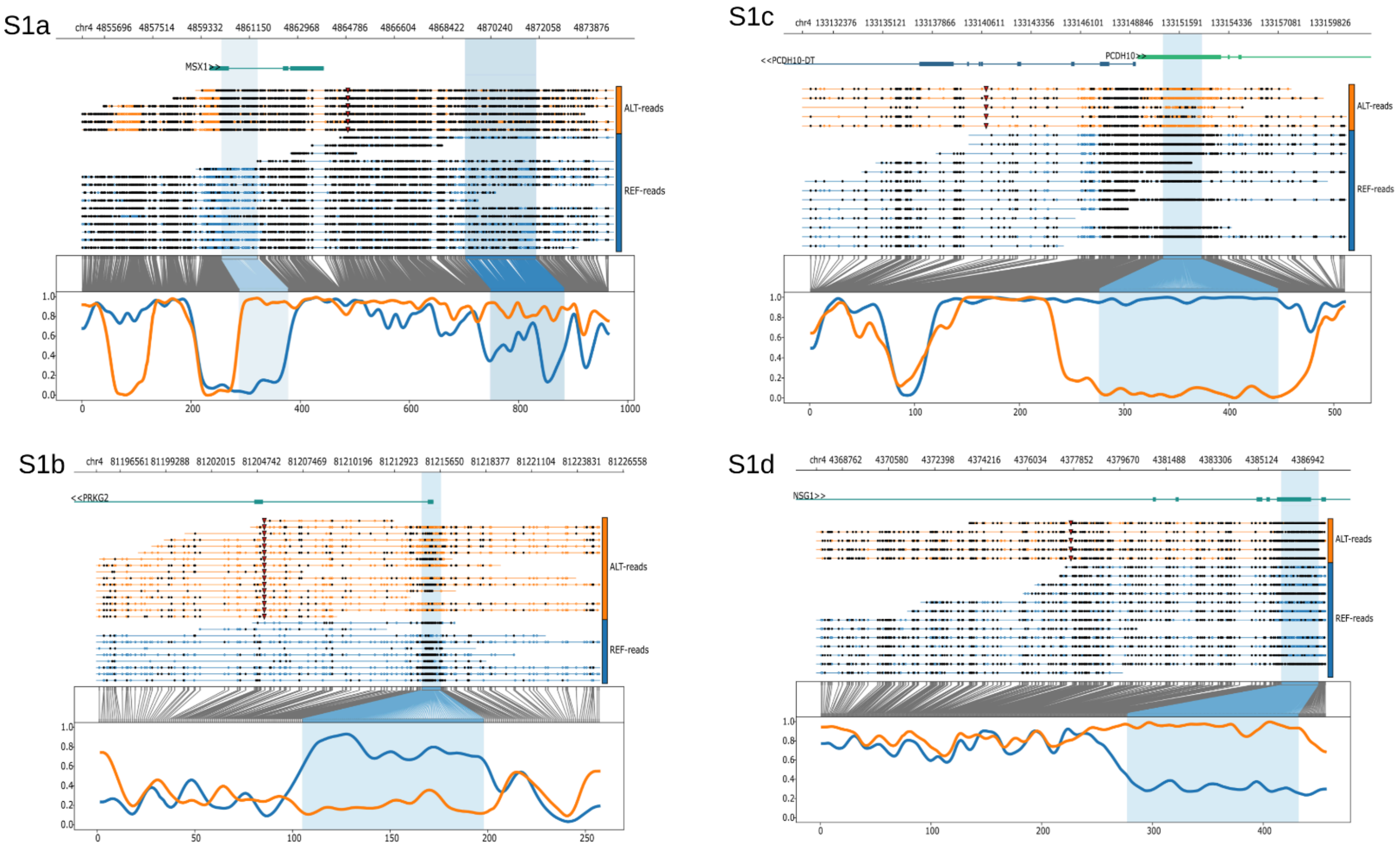


Supplementary Figure 1: Selected examples from the top 10 positive variant-methylation predictions from chr4 phased ATCC variants using unduliner. The identified differentially methylated regions are highlighted, and the triangular marks on the ALT-read groups (orange) on each plot represent the variants.

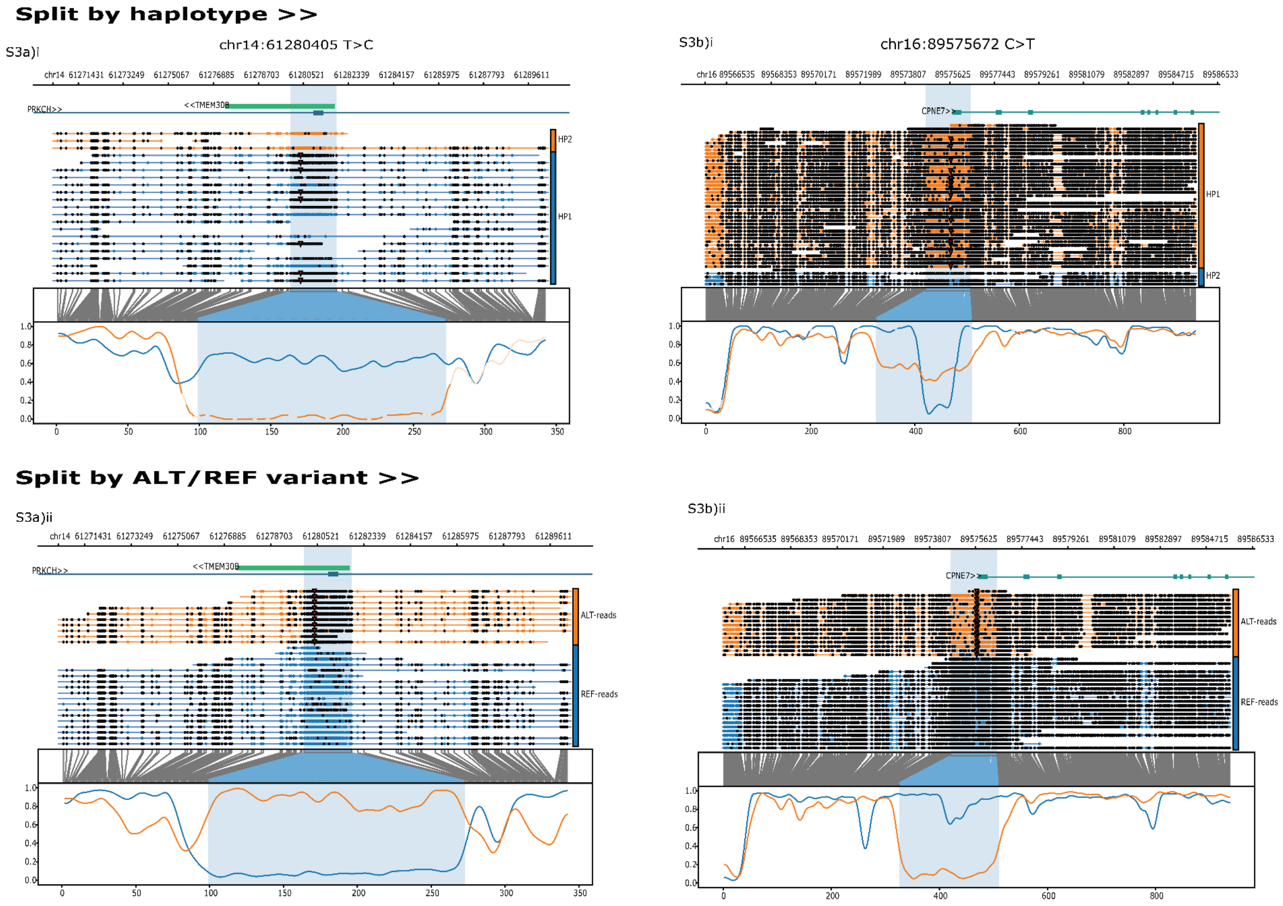


Supplementary Figure 2: Two examples of amplified variant-associated methylation differences (highlighted) in the ATCC subline compared to the haplotype. The top row shows methylation at the haplotype level with some differential methylation in the highlighted region, and the bottom row shows methylation at the variant level with increased differential methylation. Each column represents a named variant marked by triangles on each read. In all cases at the bottom row, the orange represents the reads supporting the ALT base and the blue represents the REF base.

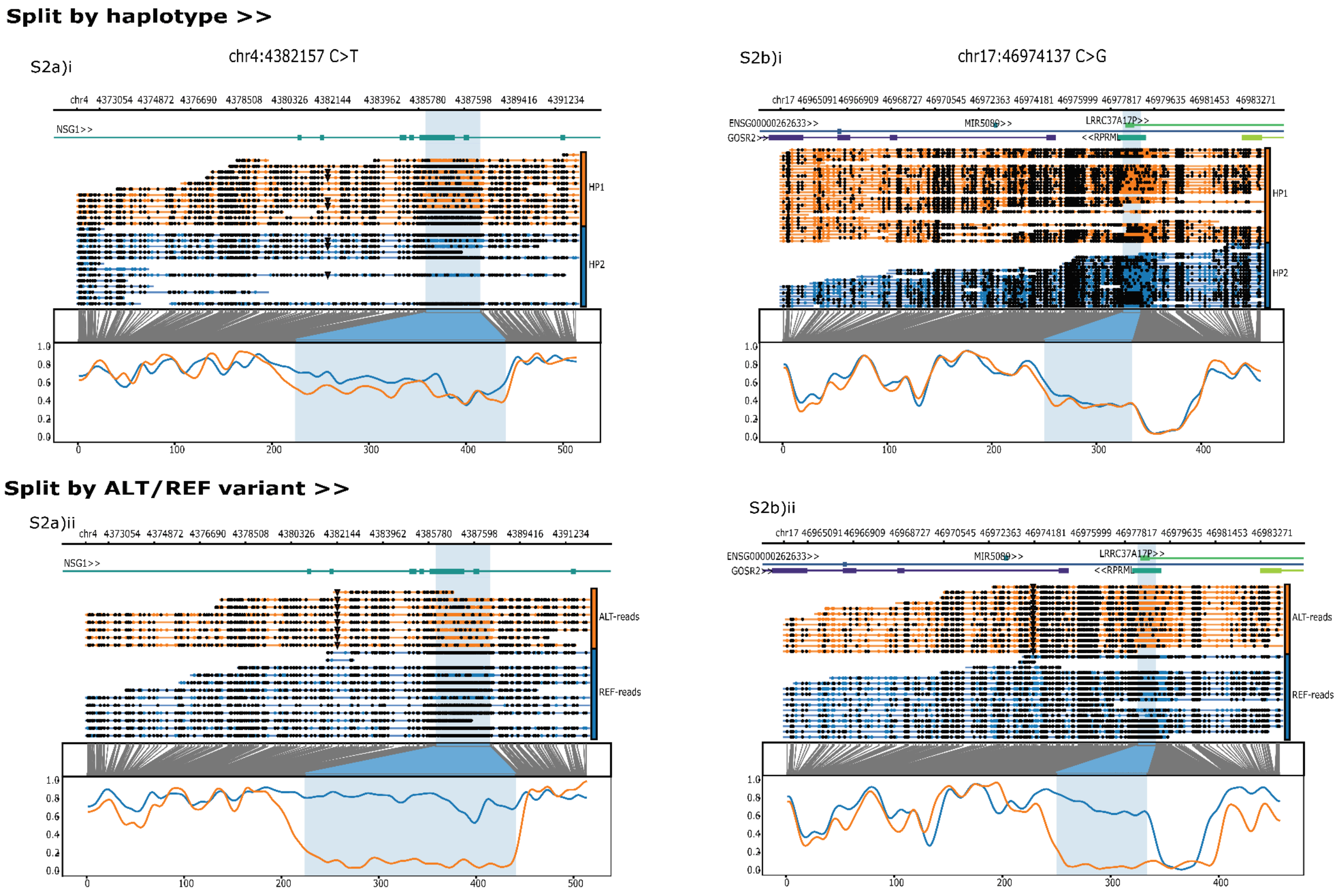


Supplementary Figure 3: Two examples of induced variant-associated methylation differences (highlighted) in the ATCC subline. The top row shows methylation at the haplotype level with no differential methylation at the highlighted regions, and the bottom row shows methylation at the variant level with differential methylation. Each column represents a named variant marked by triangles on each read. In all cases at the bottom row, the orange represents the reads supporting the ALT base, and the blue represents the REF base.

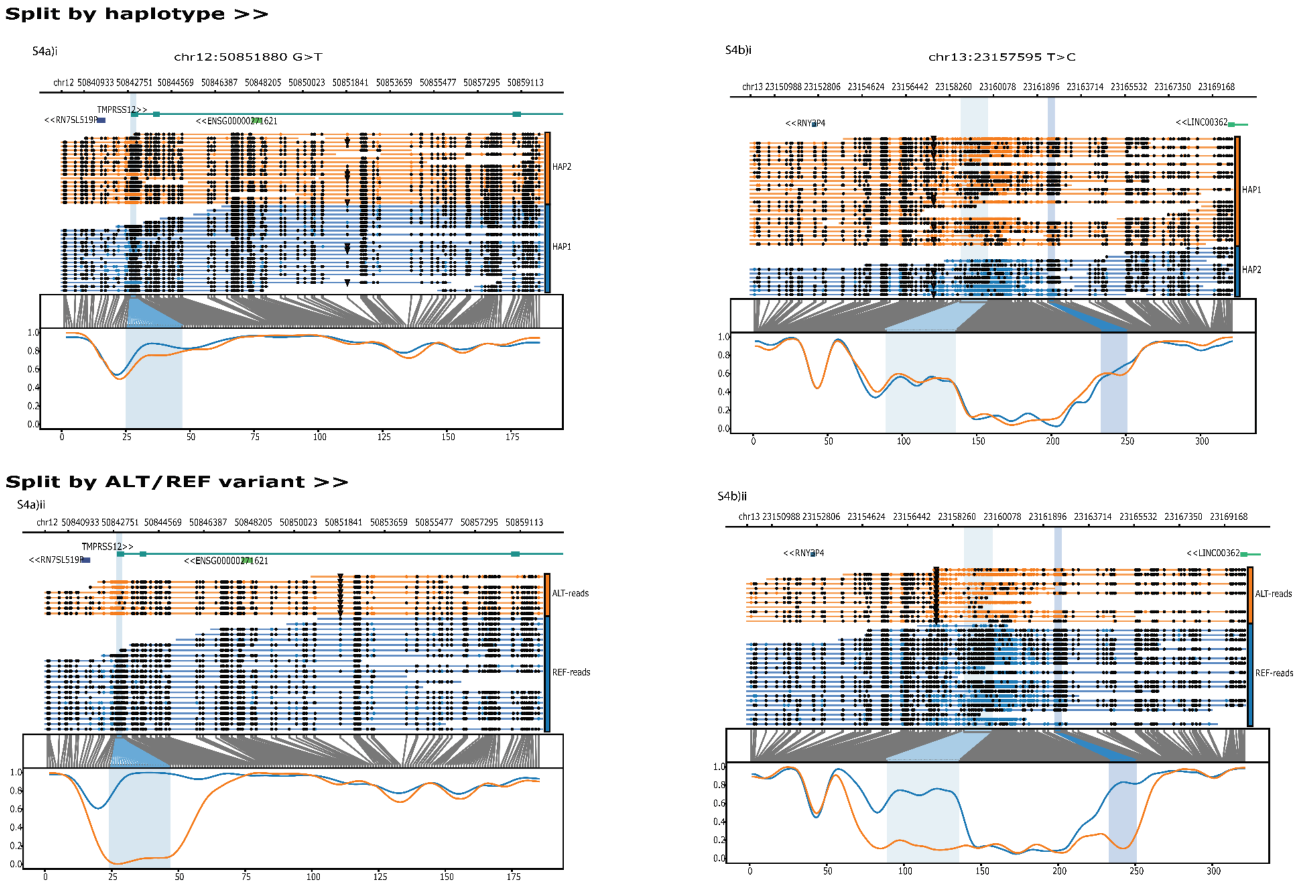


Supplementary Figure 4: Two examples of induced variant-associated methylation differences (highlighted) in the ECACC subline. The top row shows methylation at the haplotype level with no differential methylation at the highlighted regions, and the bottom row shows methylation at the variant level with differential methylation. Each column represents a named variant marked by triangles on each read.  In all cases at the bottom row, the orange represents the reads supporting the ALT base and the blue represents the REF base.

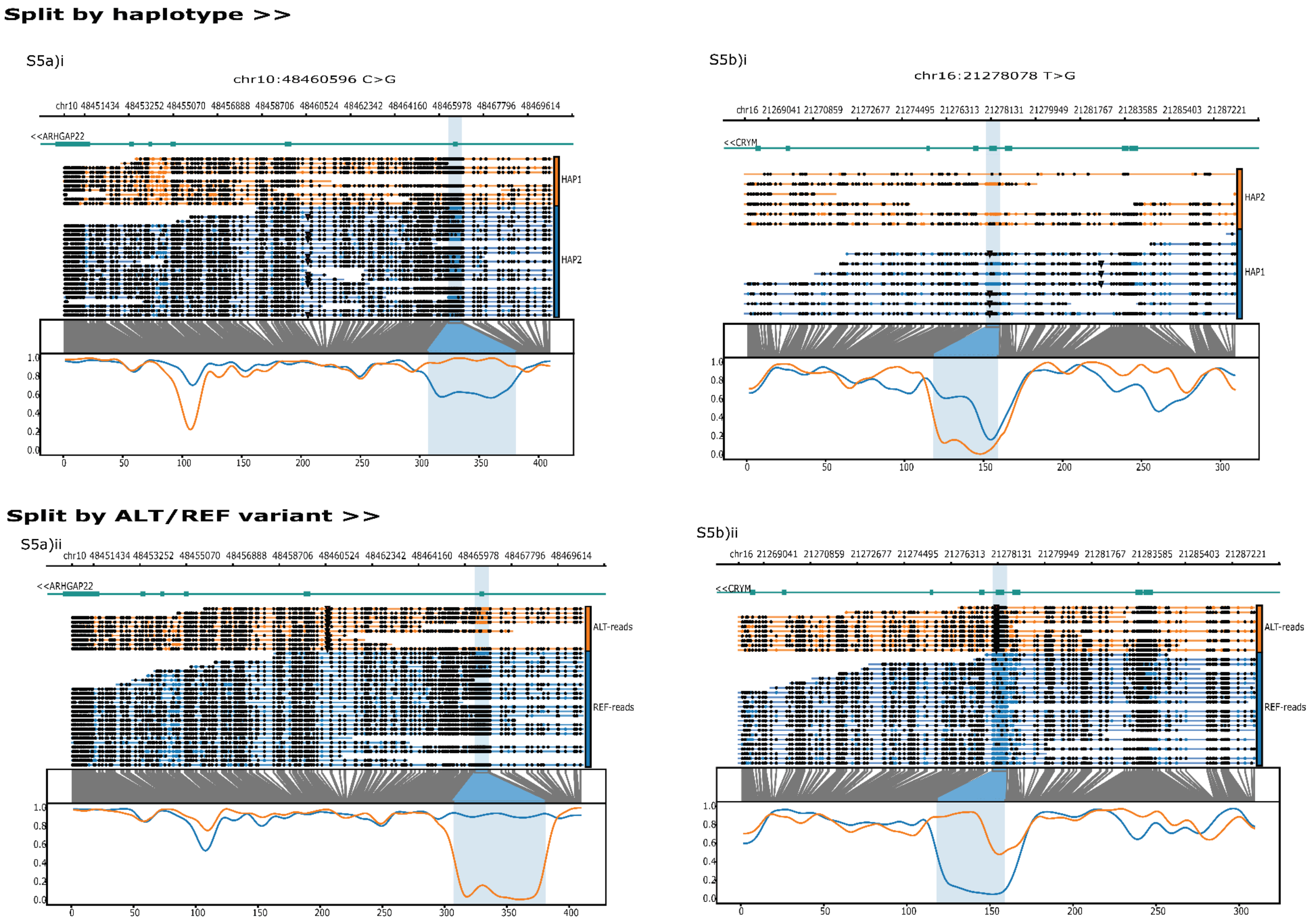


Supplementary Figure 5: Two examples of amplified variant-associated methylation differences (highlighted) in the ECACC subline. The top row shows methylation at the haplotype level with some differential methylation in the highlighted region, and the bottom row shows methylation at the variant level with increased differential methylation. Each column represents a named variant marked by triangles on each read. In all cases at the bottom row, the orange represents the reads supporting the ALT base, and the blue represents the REF base.

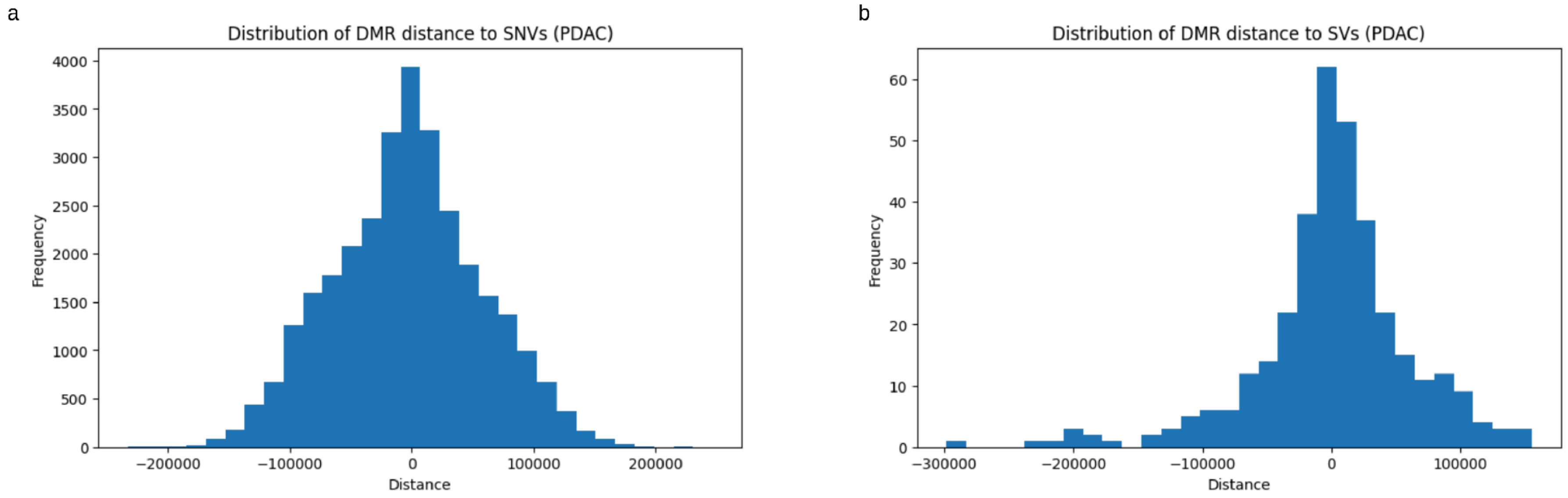


Supplementary Figure 6: The distribution of genomic range between variants (a: SNVs; b: SVs) and DMRs in the PDAC sample, where distance (x-axis) represents the difference between the variant position and end of the DMR identified by unduliner.

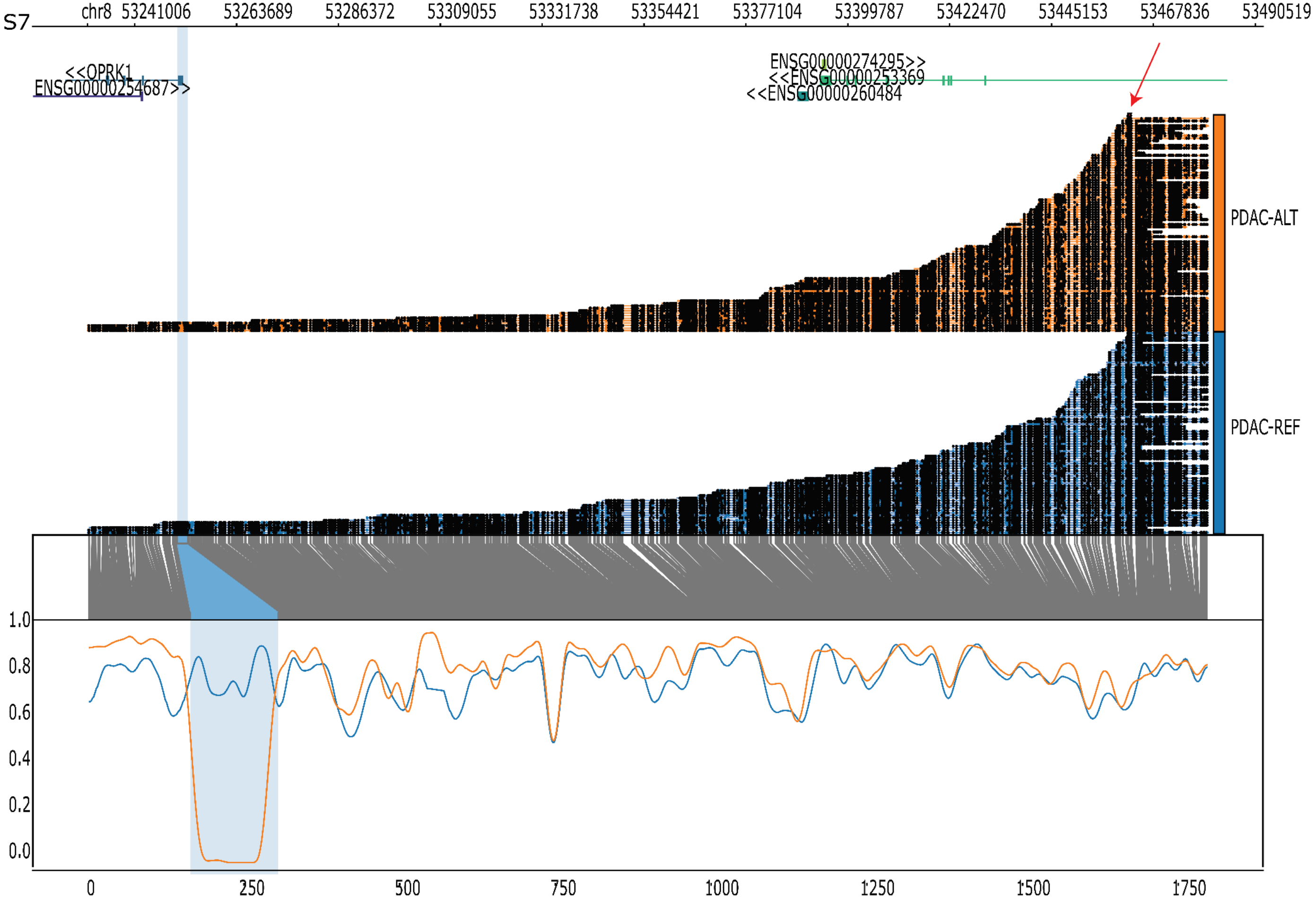


Supplementary Figure 7: An SNV (chr8:53462518 C>A) associated with differential methylation on a region 209kbp upstream (53250482-53252804). The red arrow points to the variant, which is indicated by triangle markers on all affected reads (orange).

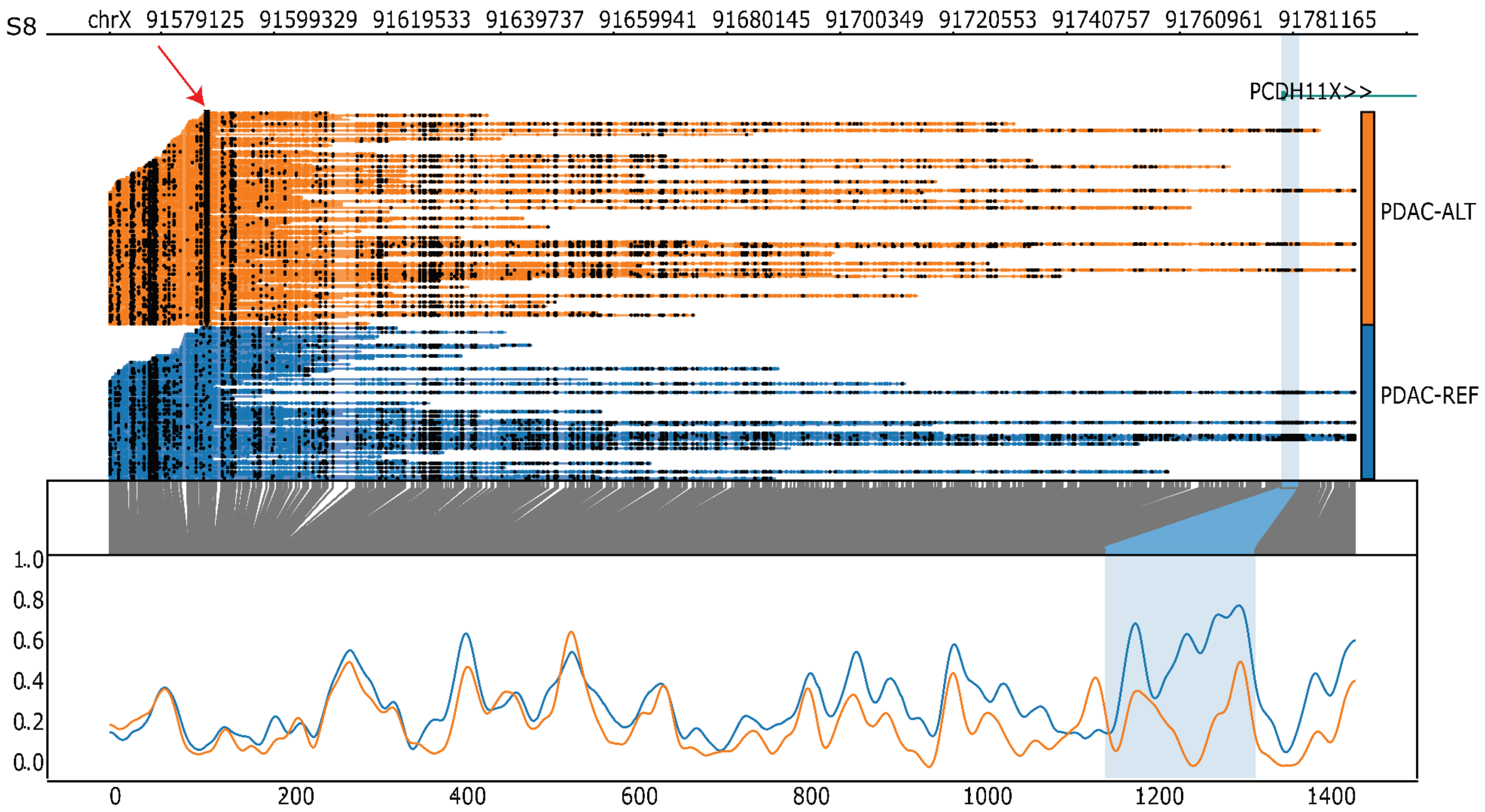


Supplementary Figure 8: An SNV (chrX:91587325 G>A) associated with differential methylation on a region ~194kbp downstream (91779002-91782252). The red arrow points to the variant, which is indicated by triangle markers on all affected reads (orange).

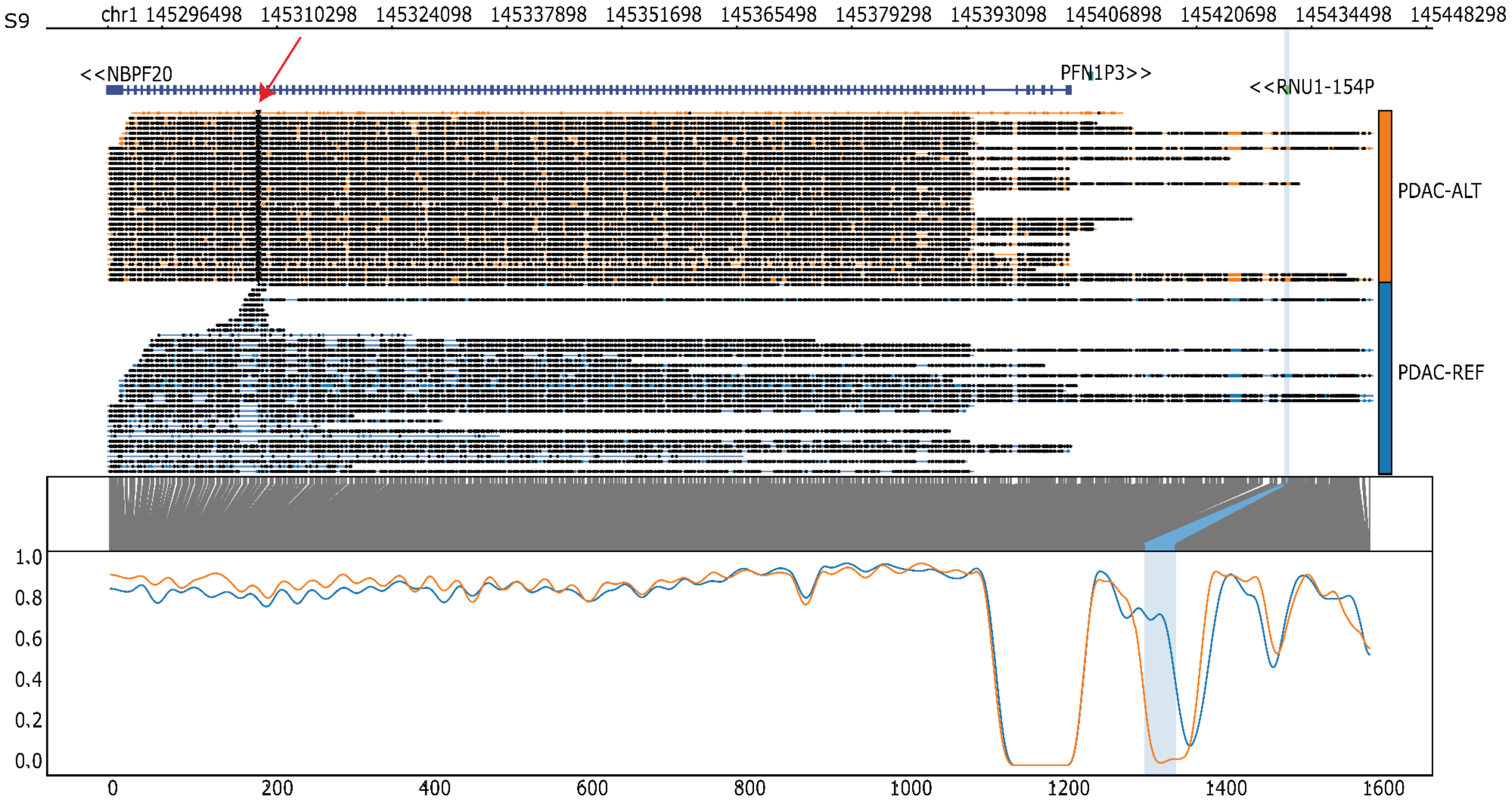


Supplementary Figure 9: An ~1.5kbp insertion on chr1 associated with differential methylation at a region ~123kn downstream (145431241-145431804). The red arrow points to the variant, which is indicated by triangle markers on all affected reads (orange).

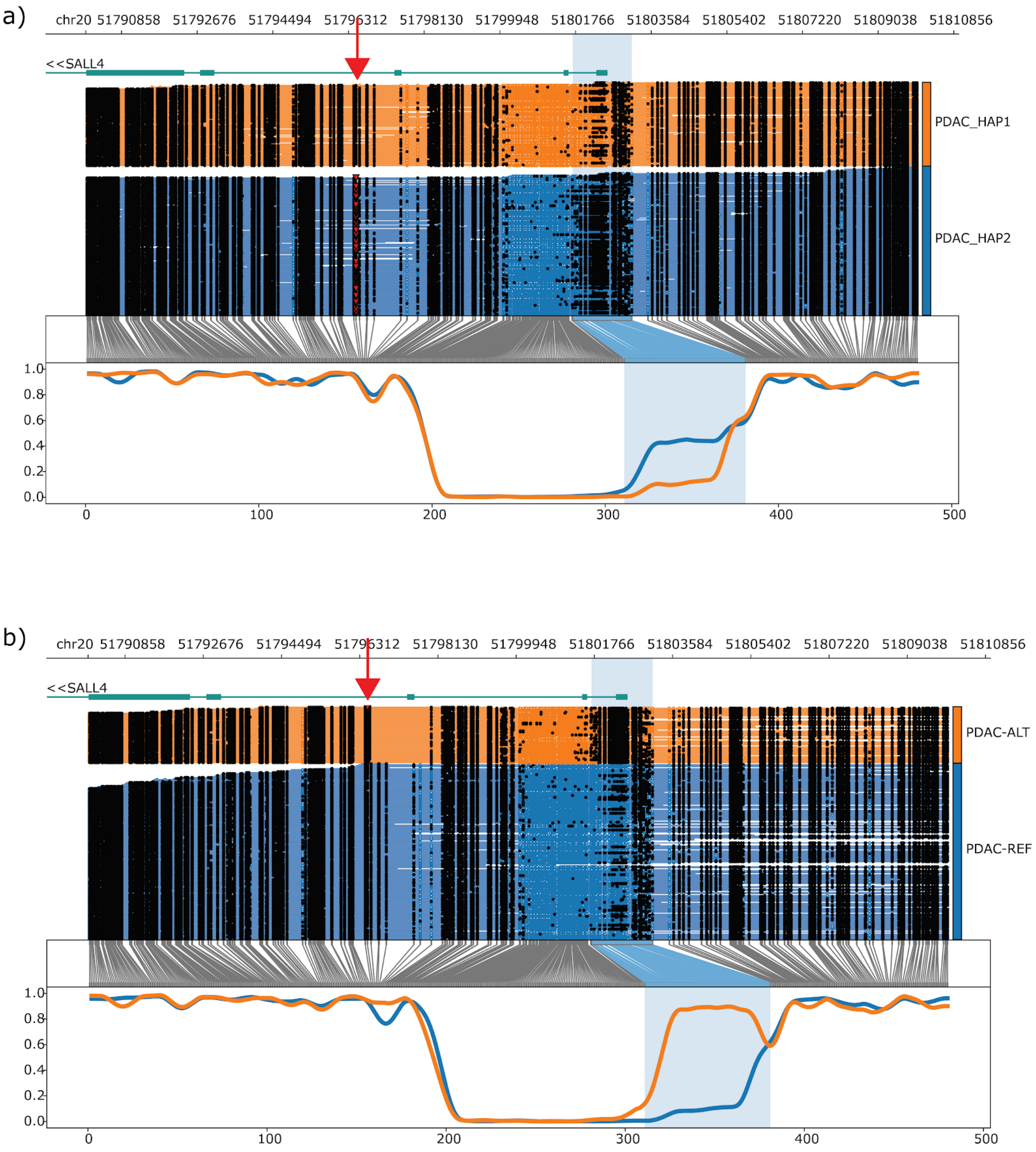


Supplementary Figure 10: Haplotype-based (a) and variant-based (b) grouping of sequence reads. A 1368-bp deletion in the SALL4 gene is limited to haplotype 2 (a) of the PDAC genome, and there is a slight hypermethylation compared to haplotype 1. When reads are grouped by variant (b), the hypermethylation is clearly amplified in the variant group compared to the reference group. The red arrow indicates the variant location.

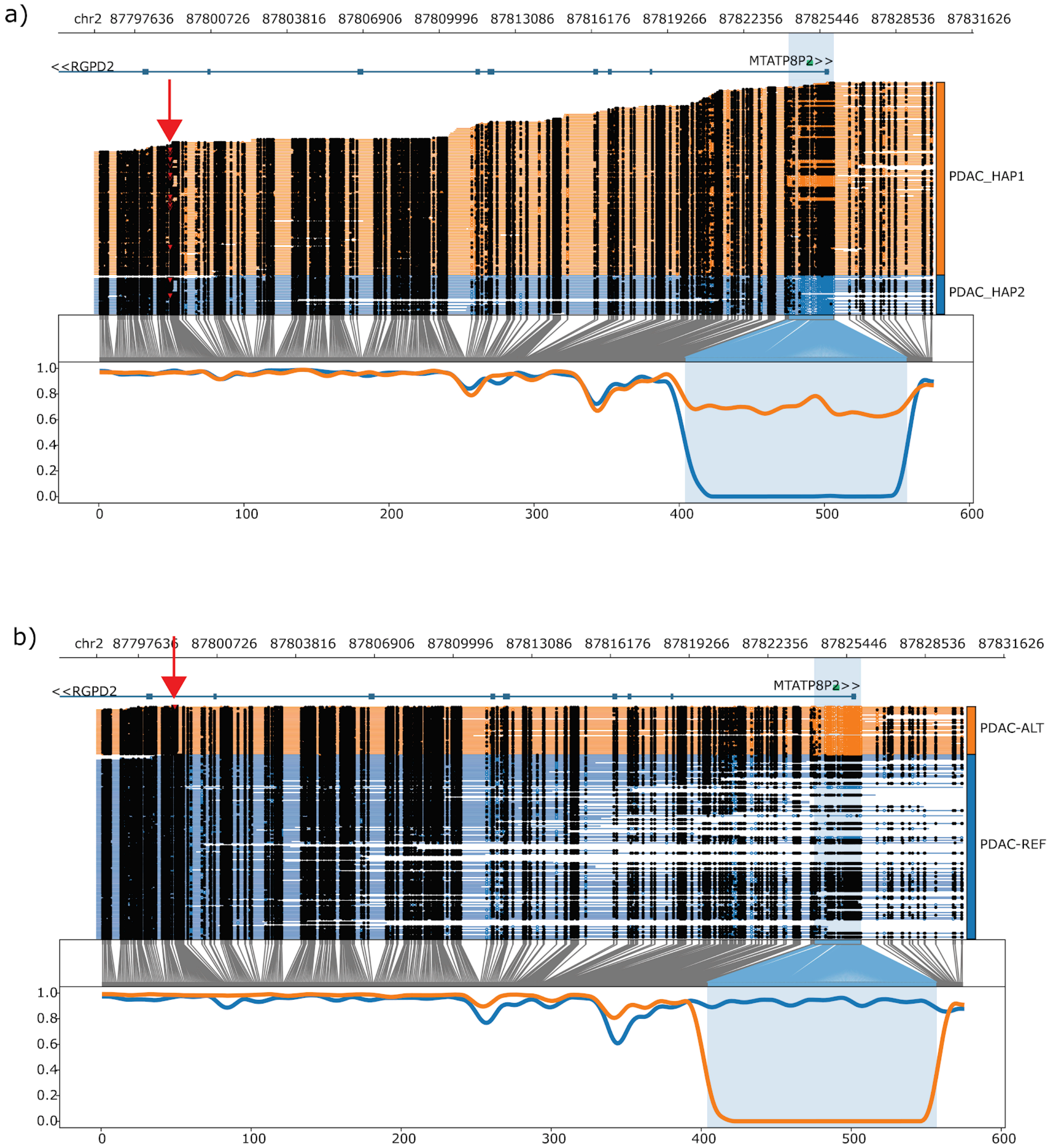


Supplementary Figure 11: Haplotype-based (a) and variant-based (b) grouping of sequence reads. A 309-bp deletion in the RGPD2 gene is present in both haplotypes (a) of the PDAC genome and exhibits haplotype-level differential methylation in an upstream region. This differential methylation becomes more clearly defined when reads are grouped by variant (b), with the variant group being completely hypomethylated compared to the reference group. The red arrow indicates the variant location.

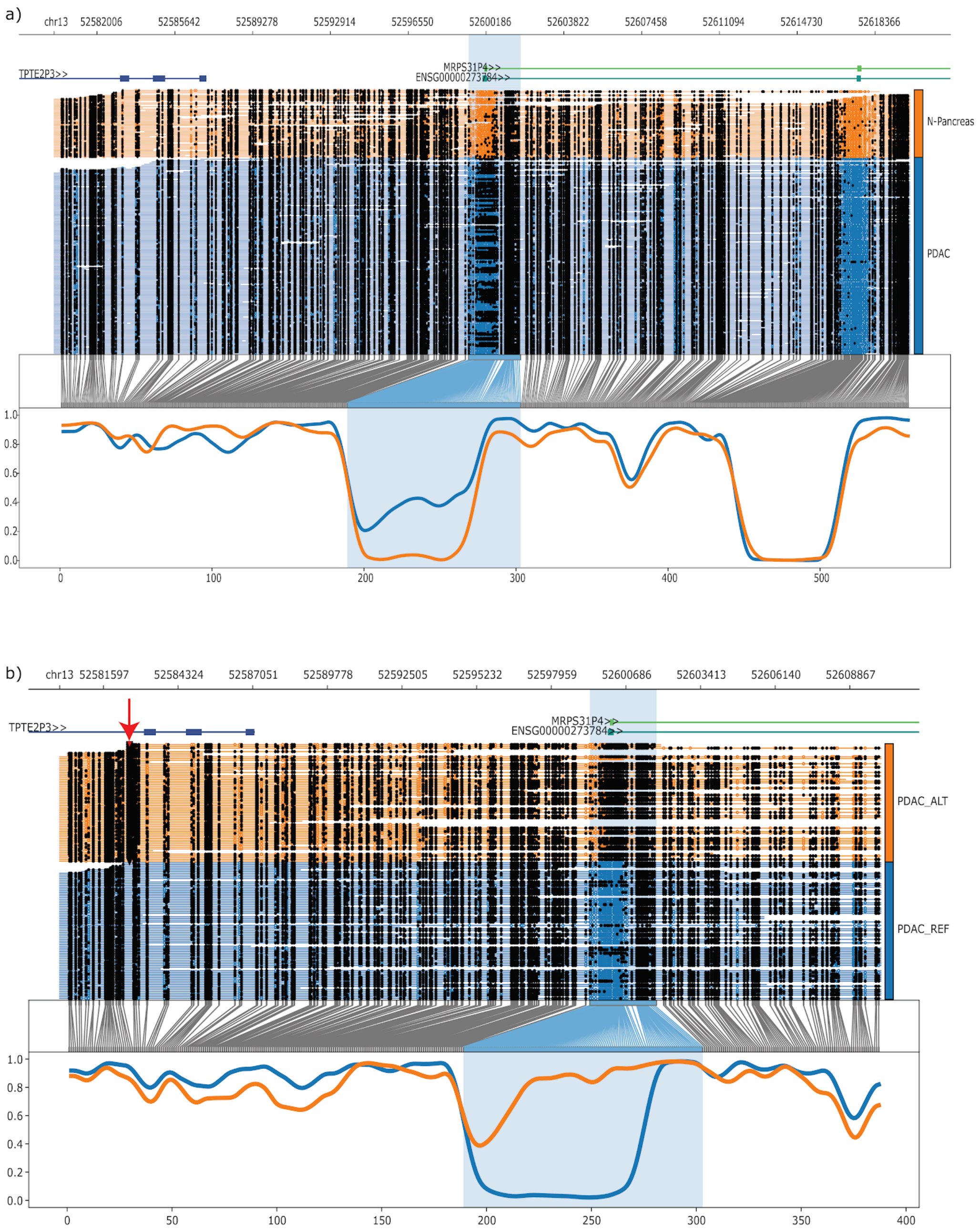


Supplementary Figure 12: An intronic non-coding variant (chr13:52582552:G>T) (b) is associated with tumour-normal (a) differential methylation overlapping enhancers and promoters. In (a), the tumour genome (blue) is hypermethylated compared to the matched normal, and in (b), the variant read group accounts for most of the observed hypermethylation. The red arrow indicates the variant location.

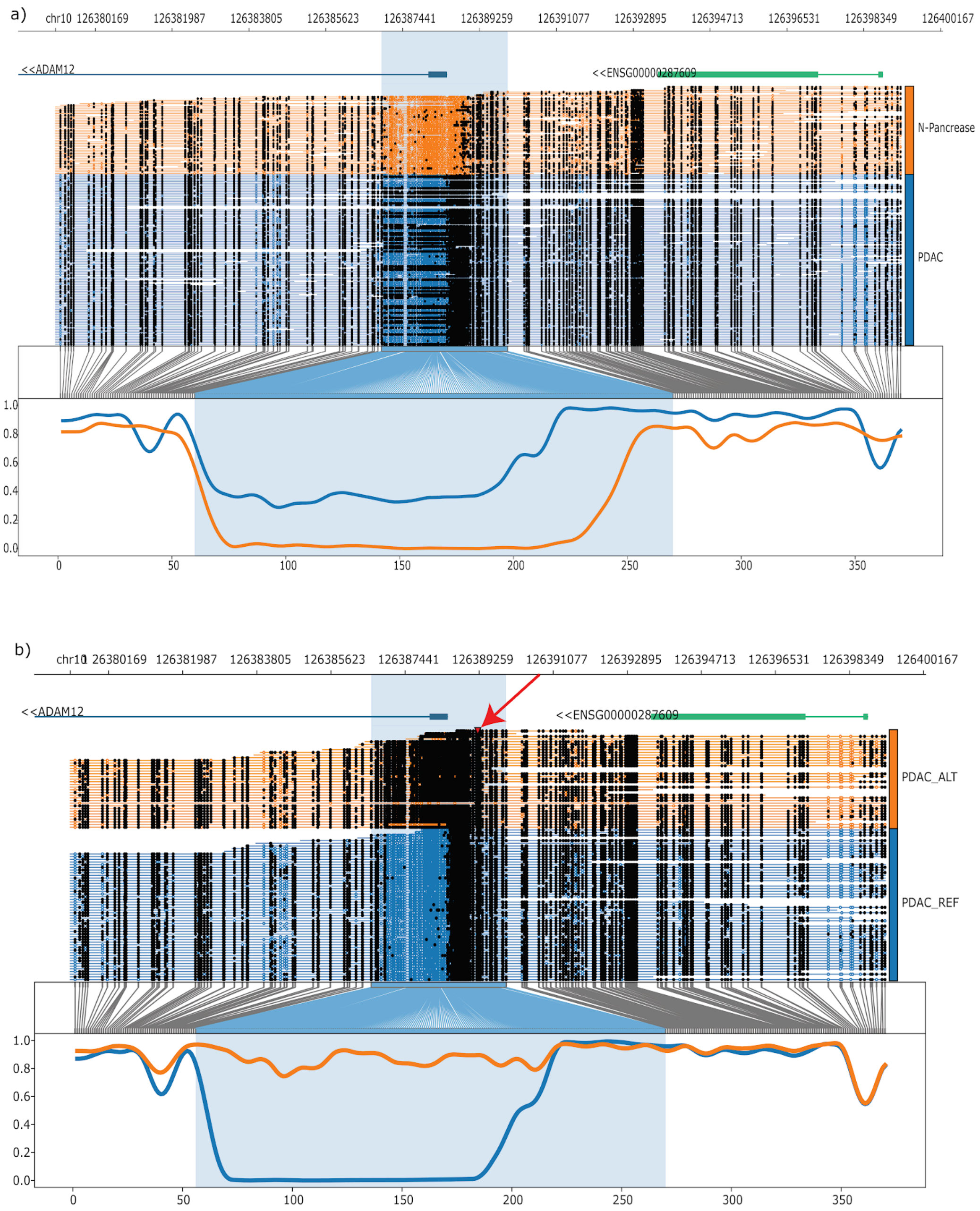


Supplementary Figure 13: A modifier variant (chr10:126389228:G>C) is associated with tumour-normal differential methylation overlapping enhancer and promoter regions (a). In (a), the tumour genome (blue) is hypermethylated compared to the normal, and in (b), the variant group is completely hypermethylated compared to the reference group. The red arrow indicates the variant location.

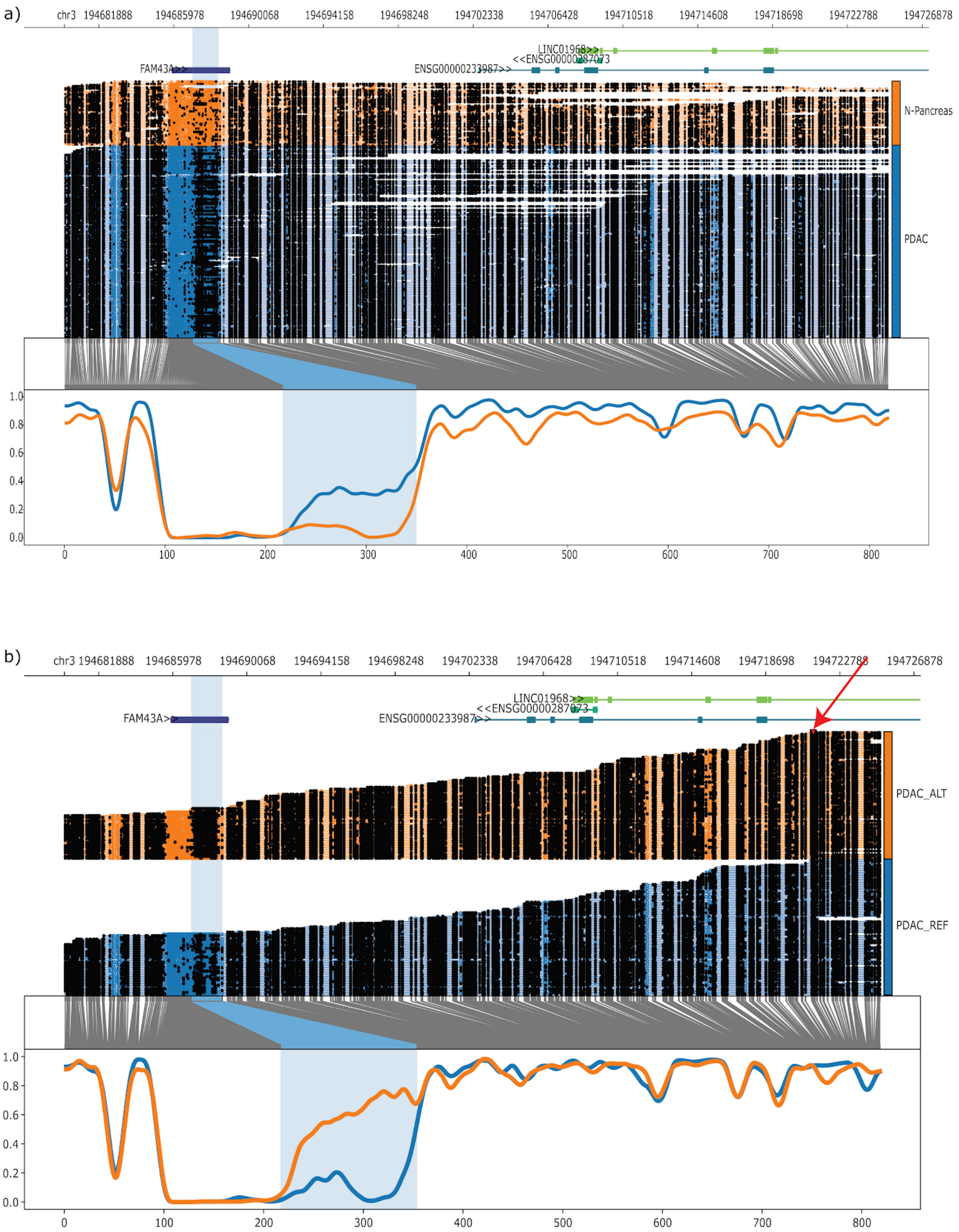


Supplementary Figure 14: An intronic variant (chr3:194721324:C>T) on a long non-coding RNA is associated with tumour-normal differential methylation (a) overlapping an enhancer. The tumour genome has a higher methylation compared to the matched normal at the highlighted region (a), and the variant reads account for a higher methylation in the tumour genome (b). The red arrow indicates the variant location.

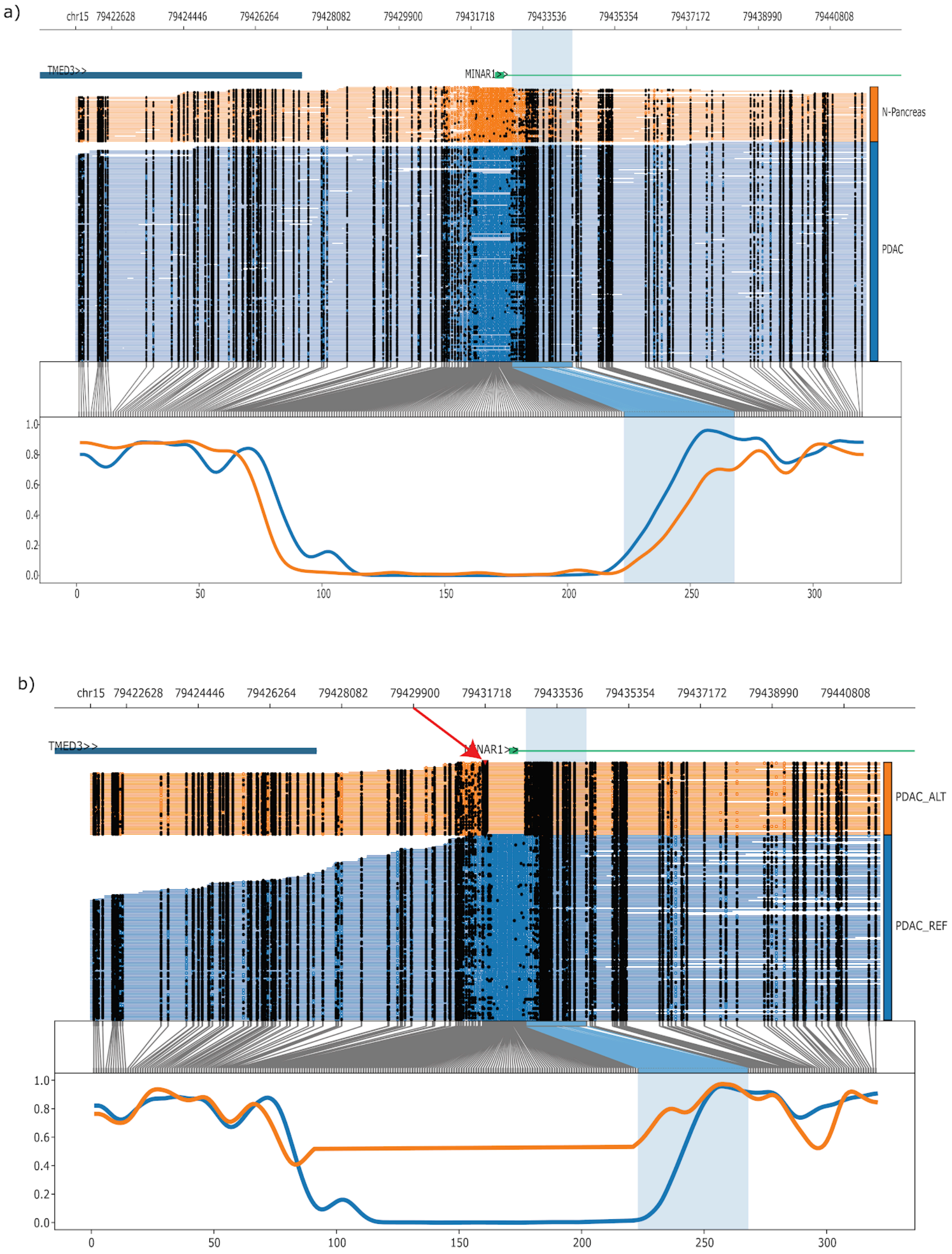


Supplementary Figure 15: A 1009-bp deletion is associated with methylation change at a region with identified tumour-normal differential methylation. The variant reads (b) are heavily methylated at the affected region compared to the reference reads. The red arrow indicates the variant location.

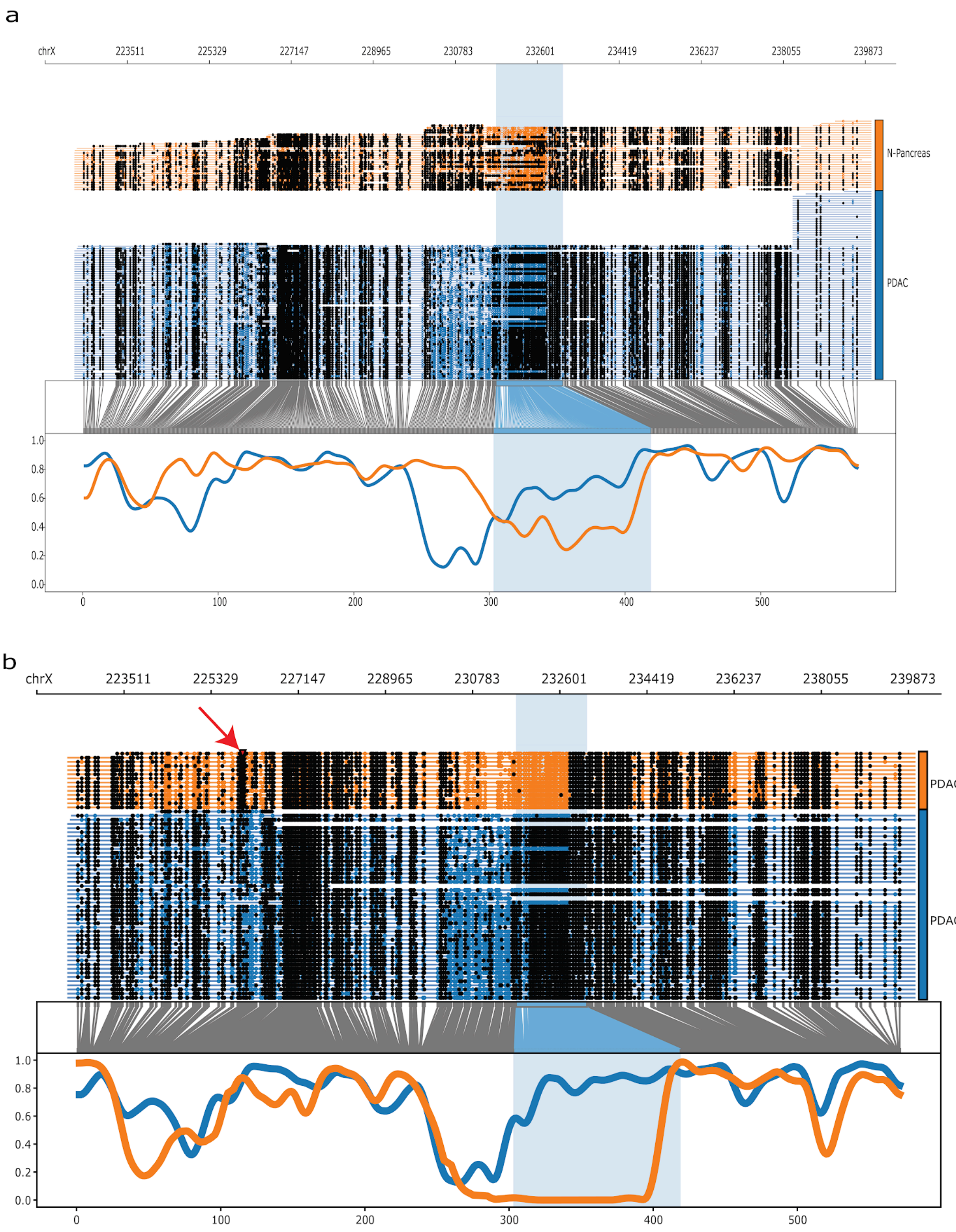


Supplementary Figure 16: A 64-bp deletion is associated with methylation change at the highlighted region, where the variant group is completely hypomethylated compared to the reference group (b). The same region is identified as differentially methylated in the tumour-normal comparison. The red arrow indicates the variant location.

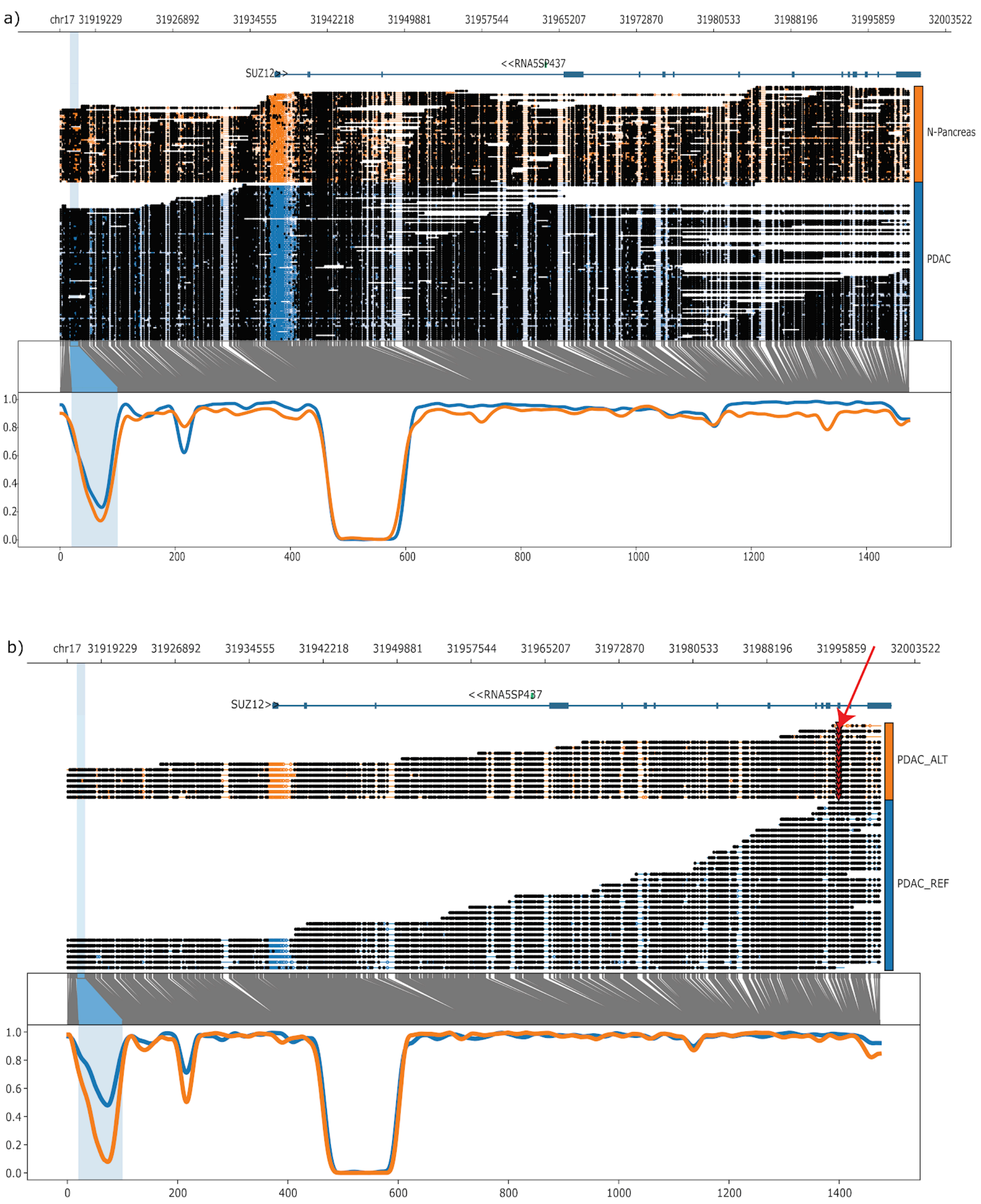


Supplementary Figure 17: No methylation difference was observed in tumour-normal comparison (a) at the highlighted region. Unduliner identified a stop-gained mutation in SUZ12 (chr17:31995631:A>T) to be associated with intratumour differential methylation. The red arrow indicates the variant location.

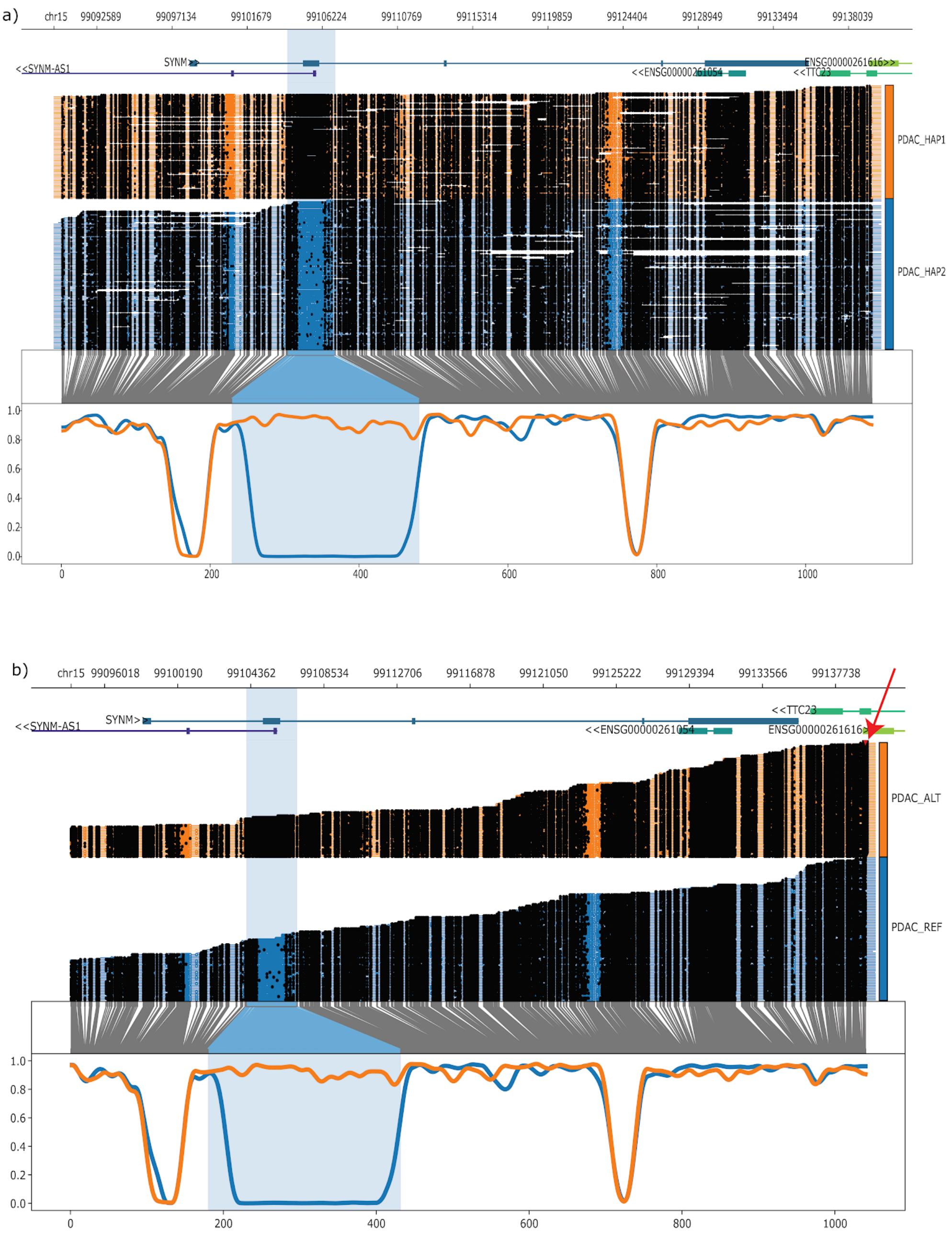


Supplementary Figure 18: Allele-specific methylation (a) is observed in the PDAC genome overlapping the SYNM/SYNM-AS1 region. A missense variant (chr15:99139442:C>T, indicated by the red arrow in b) in TTC23 was found to be associated with differential methylation in the same region.

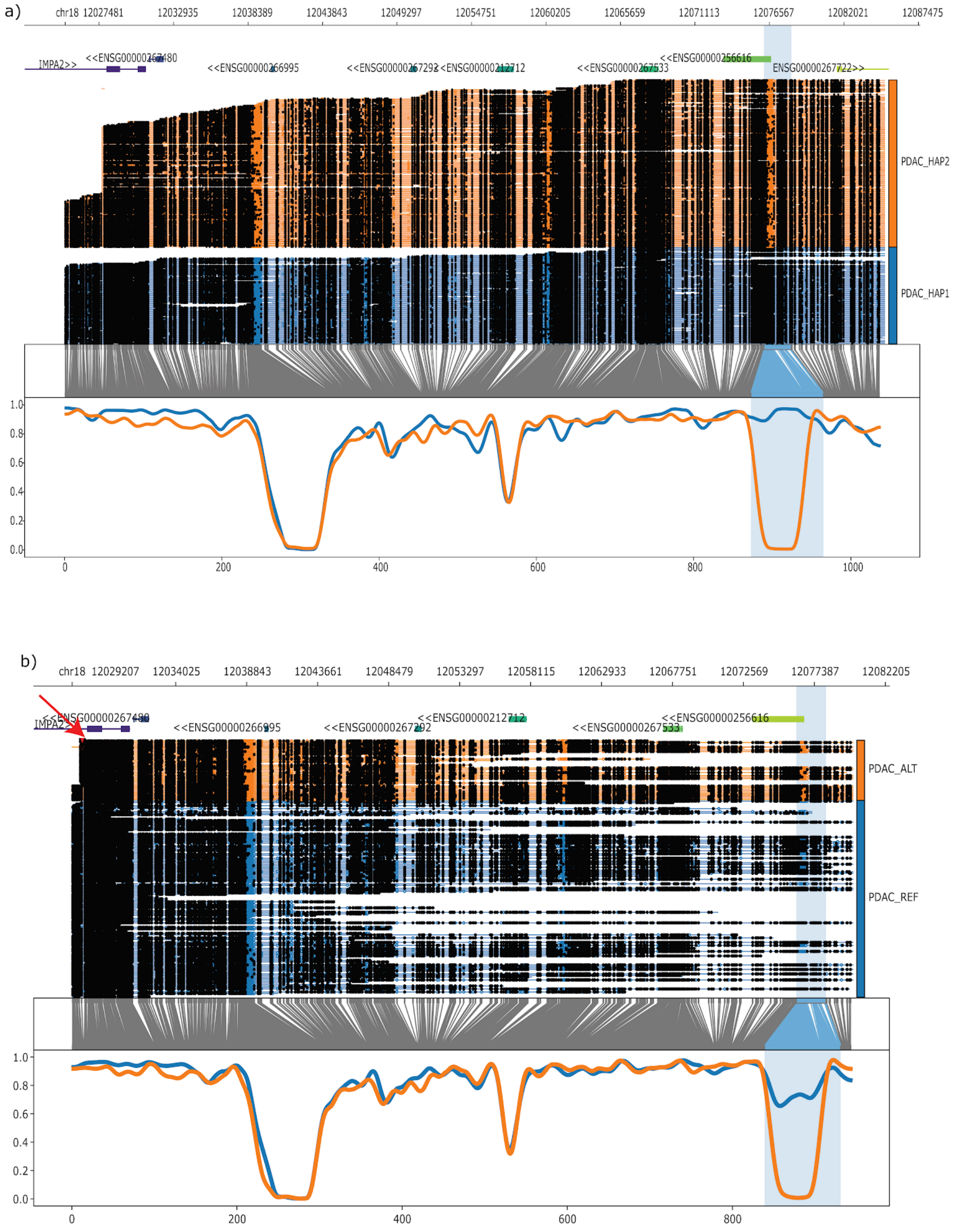


Supplementary Figure 19: Allele-specific differential methylation is present in the PDAC genome at the highlighted region (a). A duplication event (indicated by the red arrow in b) is found to be associated with differential methylation in the same region.
